# Deciphering the differential physiological and molecular requirements for conidial anastomosis tube fusion and germ tube formation in *Colletotrichum gloeosporioides*

**DOI:** 10.1101/2021.01.22.427748

**Authors:** Nikita Mehta, Ravindra Patil, Abhishek Baghela

## Abstract

The conidia of a hemibiotrophic fungus *Colletotrichum gloeosporioides* can conventionally form germ tube (GT) and develop in to a fungal colony, while under certain conditions, they tend to get connected with each other through conidial anastomosis tube (CAT) so as to share the nutrients. CAT fusion is believed to be responsible for generation of genetic variations in few asexual fungi, which appears problematic for effective fungal disease management. The physiological and molecular mechanism underlying the GT versus CAT formation remained unexplored. In the present study, we have deciphered the decision switch responsible for GT formation versus CAT fusion in *C. gloeosporioides*. GT formation occurred at high frequency in the presence of nutrients, while CAT fusion was found to be higher in absence of nutrients. Younger conidia were found to form GT efficiently, whilst older conidia preferentially formed CAT. Whole transcriptome analysis of GT and CAT fusion revealed differential molecular requirements for these two processes. We identified 11050 and 9786 differentially expressed genes (DEGs) in GT and CAT, respectively. A total 1567 effector candidates were identified, of them 103 and 101 were uniquely secreted during GT and CAT formation respectively. Genes coding for cell wall degrading enzymes, germination, hyphal growth, host-fungus interaction and virulence were up-regulated during GT formation. Whilst, genes involved in stress response, cell wall remodelling, membrane transport, cytoskeleton, cell cycle, and cell rescue were highly up-regulated during CAT fusion. To conclude, the GT and CAT fusion were found to be mutually exclusive processes, requiring differential physiological conditions and sets of DEGs in *C. gloeosporioides*. This will help to understand the basic CAT biology in the genus *Colletotrichum*.

## 1. Introduction

The asexual spore or conidium is important in life cycle of many fungi because it is the key for dispersal and serves as a ‘safe house’ for the fungi in unfavorable environmental conditions. Under favorable conditions, the conidia germinate to form hyphae and finally fully grown fungal colony. The mechanics of conidial germination are diverse and varies between different species of fungi (Osherov and May, 2001). In general, the first morphological change in conidial germination is isotropic growth also called swelling, during this stage diameter of the spore increases two-fold or more; then the next stage is polarized growth that results in the formation of a germ tube (GT) that extends and successively branches to establish the fungal colony (van Leeuwen *et al*., 2013). However, under certain conditions e.g., starvation, a conidium chooses to form specialized hypha, called conidial anastomosis tubes (CATs) instead of forming GTs. The CAT fusion between conidia result in an interconnected germling network, allowing genetic exchange, sharing of nutrients, water and cell organelles between conidia (Read *et al*., 2009), which supposedly improve the chances of survival in adverse environment.

The CATs have shown to be morphologically and physiologically distinct from GTs in *Colletotrichum lindemuthianum, Neurospora crassa* and *Fusarium oxysporum* (Kurian *et al*., 2018; Roca *et al*., 2005a, 2003). In general CATs are short, thin, usually unbranched compare to GTs. CATs grow towards each other, whilst GTs avoids each other (Roca *et al*., 2005c). It has also been shown that GT formation and CAT fusion are antagonistic in *Colletotrichum fructicola, Colletotrichum nymphaeae* and *Colletotrichum theobromicola* (Gonçalves *et al*., 2016). Interestingly, it has been shown that older conidia were optimal for CAT fusion whilst younger conidia were optimal for GT formation in *C. lindemuthianum;* and in an inter-specific CAT fusion between *C. gloeosporioides* and *C. siamense* (Ishikawa *et al*., 2010; Mehta and Baghela 2020); however, there is no such differential age requirements for CAT fusion versus GT formation in *N. crassa* and *F. oxysporum* (Kurian *et al*., 2018; Roca *et al*., 2005a; Shahi *et al*., 2016). Therefore, it is not clear that how conidial age can influence their propensity to undergo CAT fusion versus GT formation. It has been suggested that CAT fusion might play a vital role in life cycle of fungi by improving the chances of colony establishment during nutrient starvation (Roca *et al*., 2005c); however, there are contrasting reports on the relationship between CAT induction and nutrient availability e.g. CAT fusion in *C. lindemuthianum* is inhibited by nutrients and only occurs in water (Roca *et al*., 2003), while in *N. crassa* and *F. oxysporum*, CAT induction requires limited amount of nutrients to occur (Kurian *et al*., 2018; Roca *et al*., 2005a). Therefore, the effect of conidial age and availability of nutrients on CAT fusion and GT formation is poorly understood. At the gene expression level, germination of conidia in *Aspergillus niger* and *C. fructicola* has been studied using transcriptome analysis (Leeuwen *et al*., 2013; Zhang *et al*., 2018); however, whether the differential gene expression (DGE) profiles of GT remains same in case of *C. gloeosporioides* is remain to understand. Further, the transcriptome analysis of CAT fusion has never been reported in general in any fungi and in *C. gloeosporioides* in particular.

The above-mentioned points highlight a fact that the underlying physiological and molecular requirements for GT formation and CAT fusion are still not fully understood. It has been shown previously that the inter-specific CAT fusion between *C. lindemuthianum* and *C. gossypii*, resulted in some hybrids, which exhibited phenotypic variations and one hybrid turned out to be most pathogenic isolate so far reported (Roca *et al*., 2004). We have also shown that Indian *C. gloeosporioides* isolates showed high levels of genetic diversity (Mehta *et al*., 2017). Furthermore, we have shown that an inter-specific CAT fusion between *C. gloeosporioides* and *C. siamense* generated genetic and phenotypic diversity in these fungal pathogens (Mehta and Baghela, 2020). Pathogen populations with higher genetic variations tend to show increased mean fitness, resilience, expansion of host range and fungicides resistance, therefore, it is imperative to understand the basic biology of CAT fusion and decision making of GT versus CAT, as CAT fusion has been implicated for generation of genetic variations in asexual fungi. Understanding the unique and specific molecular requirements for CAT fusion and GT formation will add to the understanding of basic biology of CAT fusion and also help to develop certain drugs targeting the CAT process, thereby minimizing the genetic variations, thereby helping in effective management of asexual fungal pathogens in agricultural practices.

In the present manuscript, we have used *C. gloeosporioides* as a proxy to understand the unique and specific molecular requirements for CAT fusion and GT formation. It also starts its life cycle from conidial germination to the formation of melanized infection structures, appressoria (De Silva *et al*., 2017). The formation of GTs, appressoria and CATs in *C. gloeosporioides* was reported on apple leaves (Araujo and Stadnik, 2013), however what are the nutritional, physiological and molecular requirements for GT formation versus CAT fusion *in-vitro* are not known. In the present work, we systematically studied the *in-vitro* dynamics, nutritional and physiological requirements of GT formation and CAT fusion in *C. gloeosporioides*. Further, a comparative transcriptome analysis of CAT fusion and GT formation was done to decipher the underlying differential molecular requirements for these two processes in *C. gloeosporioides*.

## 2. Material and method

### 2.1. In-vitro dynamics of GT formation and CAT fusion

To study whether GT formation and CAT fusion are dependent on conidial age and are they mutually exclusive or not, the GT and CAT induction assays were carried out by following a protocol develop in our lab (Mehta and Baghela, 2020). The *C. gloeosporioides* (CBS 953.97) culture was inoculated on bean pod agar medium (autoclaved French bean pods submerged in 2% water agar) and was incubated in the dark at 25°C to induce sporulation. Post-inoculation the conidia were harvested at different time points *viz*. 6, 10, 13 and 17 days and a conidial suspension of 4×10^5^ per ml were prepared in distilled water and Potato dextrose broth (PDB) for CAT and GT induction, respectively. One ml of such conidial suspensions was placed in a 24-well tissue culture plate (Tarsons, India) and incubated in dark at 25°C for different time points *viz*. 3, 6, 12, 18 and 24 h for GT formation; and 24, 48, 72 and 96 h for CAT fusion. They were then examined under a microscope (Olympus BX53 with Olympus DP73 camera, Cellsens 1.13 imaging software) using differential interference contrast (DIC) optics. The GT formation and CAT fusion was quantified as the percentage of conidia (n=300) involved in GT formation or CAT fusion (Roca *et al*., 2003). Different stages of GT formation like initiation, adhesion and elongation as well as different stages of CAT formation like induction, homing and fusion were examined at different time points.

Different numbers of conidia were tested to find out the threshold of conidial numbers required for CAT fusion and GT formation in *C. gloeosporioides*. For GT induction, different concentrations of 6 d old conidia were incubated in PDB for 18 h in dark; whilst for CAT induction; different concentrations of 17 d old conidia were incubated in dH_2_O for 72 h in dark. The numbers of conidia tested were 4×10^2^, 4×10^3^, 4×10^4^, 4×10^5^ and 4×10^6^ per ml (Mehta and Baghela, 2020).

### 2.2. Effects of availability of nutrients on GT and CAT induction

In order to study whether CAT fusion and GT formation in *C. gloeosporioides* is dependent on availability of nutrients and/or starvation stress conditions, CAT fusion (in 17 d old conidia) and GT formation (in 6 d old conidia) were induced in nutrients limiting condition (water), and in presence of nutrients like PDB, 2% Glucose. To study whether the CAT induction or GT formation in *C. gloeosporioides* is mediated through MAP kinase kinase (MEK) pathway, CAT fusion and GT formation percentage was also determined in the presence of a MEK inhibitor InSolution™ PD98059 (5μM).

### 2.3. Whole transcriptome analysis of resting conidia, GT and CAT

#### 2.3.1. RNA extraction, quantification, and qualification

For transcriptome analysis, total RNA was isolated from resting conidia, germinating conidia (GTs) and fused conidia via CAT (CATs) (3 replicates of each sample) of *C. gloeosporioides* using NucleoSpin RNA plant kit (Macherey-Nagel, Germany) according to manufacturer’s guidelines. DNAse treatment was given using the Amplification grade DNase I kit (Sigma Aldrich, USA). Isolated total RNA was quantified and qualified using Nanodrop ND-1000 spectrophotometer and Qubit fluorometer. RNA quality was also assessed using formaldehyde agarose gel electrophoresis. RNA integrity was checked using a Bioanalyzer chip and Agilent RNA TapeStation (Agilent Technologies).

#### 2.3.2. Library preparation and transcriptome sequencing

500 of total RNA from the three technical replicates of each life stage of *C. gloeosporioides* were pooled and independent library preparations were carried out for three different life stages *viz*. resting conidia, GT and CAT. Pooling of RNA samples for library preparation was done due to economic constraints. RNA-seq library preparation was performed as per Illumina-compatible NEBNext UltraTM Directional RNA Library Prep. Sequencing for 150 bp length paired-end (PE) reads was performed in an Illumina HiSeq sequencer.

#### 2.3.3. Quality control, de-novo assembly, and sequence clustering

Reads were processed for quality assessment and low-quality filtering before the assembly. The raw data generated was checked for the quality using FastQC and preprocessing of the data, which includes removing the adapter sequences and low-quality bases (<q30) was done with Cutadapt (Martin, 2011).

Processed reads were assembled using graph-based approach by rnaSPAdes program (Bankevich *et al*., 2012). rnaSPAdes is a tool for *de-novo* transcriptome assembly from RNA-Seq data and is suitable for all kind of organisms. The characteristic properties, including N50 length, average length, maximum length, and minimum length of the assembled contigs were calculated. In the second step of the assembly procedure clustering of the assembled transcripts based on sequence similarity is performed using CD-HIT-EST (Fu *et al*., 2012) with 95% similarity between the sequences, which reduces the redundancy without exclusion of sequence diversity that is used for further transcript annotation and differential expression analysis.

#### 2.3.4. Read mapping to the reference genome and differential counting

To assess the quality of the assembly, evaluation of read content approach was used. Processed reads from all three libraries were aligned back to the final assembly using Bowtie2 (Langmead and Salzberg, 2012) with end-to-end parameters. *C. gloeosporioides* Nara gc5 isolate (ftp://ftp.ncbi.nlm.nih.gov/genomes/genbank/fungi/Colletotrichum_gloeosporioides/) was used as reference genome. For each sample, the count of mapped reads was derived and normalized to RPKM (reads per kilobase of exon model per million mapped reads). The count data was further used to identify differentially expressed genes at each time point using DESeq package (Anders and Huber, 2010). Sequencing (uneven library size/depth) bias among the samples was removed by library normalization using size factor calculation in DESeq. Fold changes were determined according to the formula “Expression of treated sample / Expression of control sample”. The raw P values were adjusted for multiple testing with the procedure described by (Benjamini and Hochberg, 1995) to control FDR (false discovery rate). The regulation for each transcript was assigned on the basis of their log2fold change.

#### 2.3.5. Gene ontology enrichment, differential expression analysis and functional annotation

Assembled transcripts were functionally enriched and categorized based on blast sequence homologies and gene ontology (GO) annotations describing biological processes and molecular functions using Blast2GO software (P < 0.05), selecting the NCBI blast Fungi as taxonomy filter and default parameters (Götz *et al*., 2008). Out of 1,21,260 total 88,491 transcripts (72%) were functionally annotated against all fungal protein sequences from Uniprot Protein database. Those transcripts with more than 30% identity as cut off were taken for further analysis.

Multiple databases (Uniprot, NCBI, KEGG pathway and Pfam) were used for functional annotation of the transcripts and to determine possible roles of the annotated genes/proteins. Clustered transcripts were annotated using homology approach to assign functional annotation using BLAST tool (Altschul *et al*., 1990) against “Fungi” data from the Uniprot database containing 95,147,95 protein sequences. Transcripts were assigned with a homolog protein from other organisms, if the match was found at e-value less than e^−5^ and minimum similarity greater than 30%. Top 50 highly up-regulated transcripts of each GT and CAT were further manually annotated using Ensembl Fungi database (https://fungi.ensembl.org/Colletotrichum_gloeosporioides_cg_14_gca_000446055/Info/Index) for *C. gloeosporioides* to find out more accurate molecular and biological functions.

#### 2.3.6. Transcription factors and secreted protein analysis

Assembled transcripts were translated into protein using TransDecoder. The putative transcription factors (TFs) were predicted using the Fungal Transcription Factors Database (http://ftfd.snu.ac.kr). Transcription factor analysis was performed based on homology approach using NCBI-blast program.

For effector proteins identification, proteins with signal peptides were identified using SignalP (Petersen *et al*., 2011). Proteins with a transmembrane helix predicted using TMHMM (Emanuelsson *et al*., 2007) was excluded from the analysis. Candidate effector proteins were identified by subcellular localization prediction using TargetP (Möller *et al*., 2001) and WolfPsort (Horton *et al*., 2007). All putative proteins with a SignalP D-score = Y were considered. These proteins were then scanned for transmembrane spanning regions using TMHMM and proteins with 0 transmembrane domains (TM) were retained. Finally, proteins with location predicted as Loc = S (Petersen *et al*., 2011) using TargetP and proteins predicted as extracellular (Ext >17) using WolfPSort were retained in the final candidate effector protein dataset. The Pfam search (EMBL-EBI) was performed for functional annotation and to find out the possible biological roles of secreted proteins during GT formation and CAT fusion.

#### 2.3.7. Data Availability

RNA-seq data were submitted to the NCBI SRA database under accession number SRR12245378, SRR12245377 and SRR12245376.

### 2.4. Real time qRT-PCR validation

Validation of differential gene expression obtained from the transcriptome analysis was performed using qRT-PCR assay on total cDNA samples from the 3 life stages *viz*. resting conidia, GT and CAT of *C. gloeosporioides*. For this, 27 differentially expressed genes from resting conidia, GT and CAT detected by RNA-seq were randomly chosen for validation by qPCR. Beta-tubulin gene was used as a reference gene for normalization. Specific primer pairs were designed for these 27 selected genes using the Primer3 Software, so as to generate final amplicon sizes of each gene between 50 to 150 bp with 60°C melting temperature. All primers used in this study are listed in the supplementary Table 8. Quantitative real-time PCR (qRT-PCR) was carried out as previously described (Rudd *et al*., 2015). In brief, total RNA (1 μg) from resting conidia, GTs and CATs were reverse transcribed to cDNA with an oligo (dT) primer using SuperScript III (Invitrogen, Carlsbad, CA 92008, USA) according to the manufacturer’s instructions. A regular PCR amplification for each primer pair was performed to optimize the annealing temperature. The qRT-PCR was performed in PCRmax Eco 48 Real-Time Cycler using 50 ng of cDNA, 5 μL of SYBR™ Green PCR Master Mix (Applied Biosystems), 10 pmol of sense primers and 10 pmol of antisense primers in a final volume of 10 μL. The PCR cycle conditions were set as follows: 95°C for 30 s, followed by 40 cycles of 95°C for 10 s, 55°C for 30 s and 60°C for 15 s. At last, in order to check the amplification of single targeted amplicon, melting curves were analyzed for all the primer pairs. Expression of genes was evaluated according to their relative quantification using the 2^−ΔΔCt^ method (Schefe *et al*., 2006). Each sample was run in triplicates for all selected and reference genes. Data were analyzed using the EcoStudy software v5.2. A correlation between expression levels of DEGs by qRT-PCR and RNA seq was evaluated using log2fold expression values.

### 2.5. Statistical analyses

Data were analyzed by Analysis of Variance (ANOVA) with Tukey’s post hoc test using GraphPad Prism 5 Statistics Software, wherever applicable. Differences with a *p-value*< 0.05 were considered statistically significant. All assays were performed in triplicates and with 300 conidial numbers, wherever applicable.

## 3. Results

### 3.1. CAT fusion and GT formation are mutually exclusive in C. gloeosporioides

When CAT fusion and GT formation was studied in differently aged conidia, it was observed that young conidia (6 days) in PDB were efficient in forming GT, while older conidia (17 days) in sterile distilled water were good at forming CAT (Fig. 1A). The younger (6 days) and older (17 days) conidia in PDB do not form CAT at all, while in water they showed 3% ± 1% and 13.3% ± 2.3% CAT fusion respectively. As the age of conidia increases the GT formation percentage decreases, on the contrary, the CAT fusion percentage increases with the increasing age of conidia. The young conidia (6 days) in PDB showed 19.8% ± 1.8% GT formation and no CAT fusion, while the same aged conidia in water showed 3% ± 1% CAT fusion and no GT formation (Fig. 1A). The old conidia (17 days) when incubated in water showed 13.3% ± 2.3% CAT fusion and no GT formation, while the same aged conidia in PDB showed 5.7% ± 0.6% GT formation and no CAT fusion (Fig. 1A). In order to find out the accurate incubation time required for GT formation in 6 days old conidia, when these conidia were incubated in PDB for different time points *viz*. 3, 6, 12, 18 and 24 h, the maximum GT formation (20.1% ± 4.5%) was seen at 18 hours post incubation (hpi) (Fig. 1B). Whilst, when 17 days old conidia were incubated in distilled water for 24, 48, 72 and 96 h, the maximum CAT fusion frequency (13.5% ± 2.6%) was observed at 72 hpi (Fig. 1C). Therefore, most of the further experiments were conducted using these optimized parameters for *in-vitro* GT and CAT induction.

**Fig. 1:**
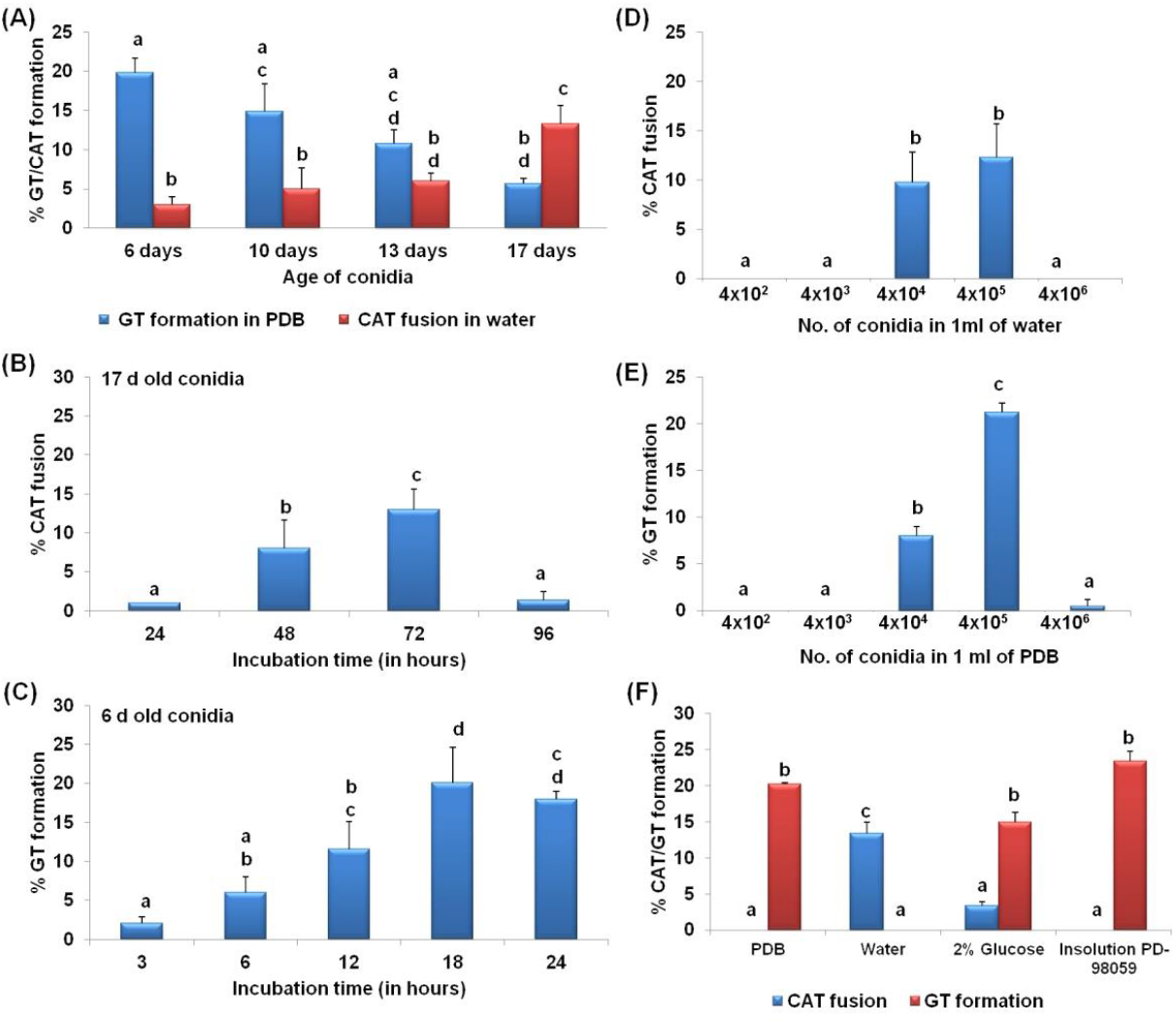
Germ tube formation and conidial anastomosis tube fusion dynamics in *C. gloeosporioides*. **A)** Percentage GT/CAT formation in different aged conidia *viz*. 6, 10, 13 and 17 days of *C. gloeosporioides*. **B)** Percentage CAT fusion in 17 days old conidia of *C. gloeosporioides* incubated for different incubation time *viz*. 24, 48, 72 and 96 h. **C)** Percentage GT formation in 6 days old conidia of *C. gloeosporioides* incubated for different incubation time *viz*. 3, 6, 12, 18 and 24 h. **D)** Percentage CAT fusion in different conidial densities *viz*. 4×10^2^, 4×10^3^, 4×10^4^, 4×10^5^ and 4×10^6^ of 17 days old conidia incubated in 1 ml of water. **E)** Percentage GT formation in different conidial densities *viz*. 4×10^2^, 4×10^3^, 4×10^4^, 4×10^5^ and 4×10^6^ of 6 days old conidia incubated in 1 ml of PDB. **F)** CAT and GT formation percentage in PDB, water, glucose, and Insolution™ PD98059 (inhibitor of MEK), CAT fusion induced in water supplemented with glucose and Insolution™ PD98059; whilst GT formation was assayed in PDB supplemented with glucose and Insolution™ PD98059. GT/CAT formation was quantified as the percentage of conidia involved in GT/CAT formation. Average from 3 replicates (n=3) and 300 conidial numbers were counted per replicate. Bar indicates standard deviation. Statistical significance of differences was analyzed by one-way ANOVA with Tukey’s multiple comparison post-hoc test (bars with the same letter are not significantly different; *p* ≤ 0.05).

Different stages of the CAT fusion in *C. gloeosporioides* consist of following stages. Initially conidia get primed for CAT fusion as a result of CAT induction (Fig. 2A). Subsequently, CATs home towards each other known as CAT homing (Fig. 2B). The CATs were fused to each other constituting the third stage i.e., CAT fusion (Fig. 2C). Later on, with increasing incubation time CAT connections expanded and formed CAT network (Fig. 2D) till 72 h of incubation. Different stages of the GT formation in *C. gloeosporioides* could be divided in to three stages namely conidial swelling and adhesion at 3 hpi (Fig. 3A-B), GT initiation includes polarized growth at 6 hpi (Fig. 3B-C), GT elongation includes hyphal growth and formation of septa 12-18 hpi (Fig. 3D-E) and finally GT network formed at and after 24 hpi (Fig. 3F).

**Fig. 2:**
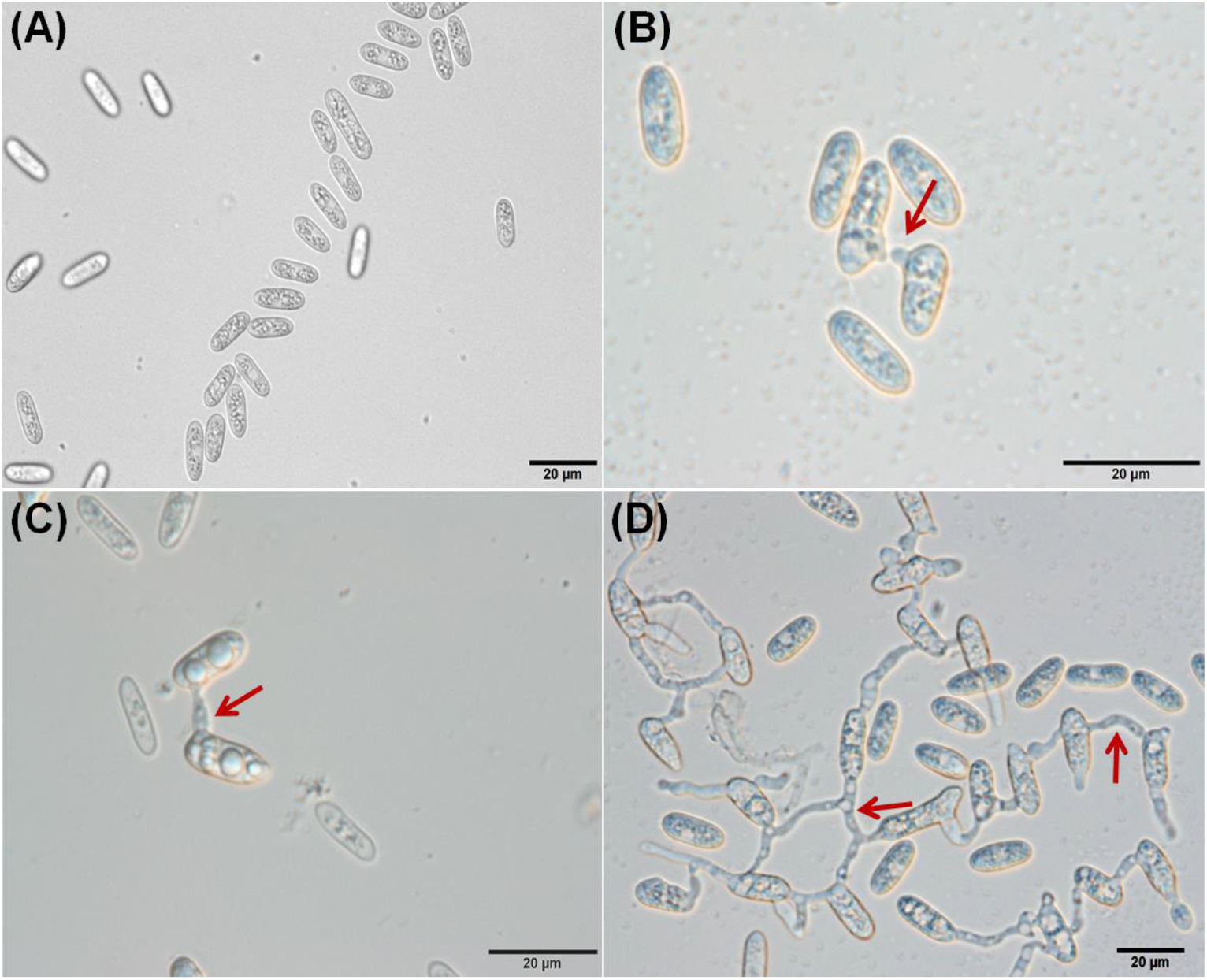
Different stages of CAT fusion in *C. gloeosporioides in-vitro*. **A)** CAT induction. **B)** CAT homing. **C)** CAT fusion. **D)** CAT network. Arrow indicates CAT fusion. Scale Bar = 20μm.

**Fig. 3:**
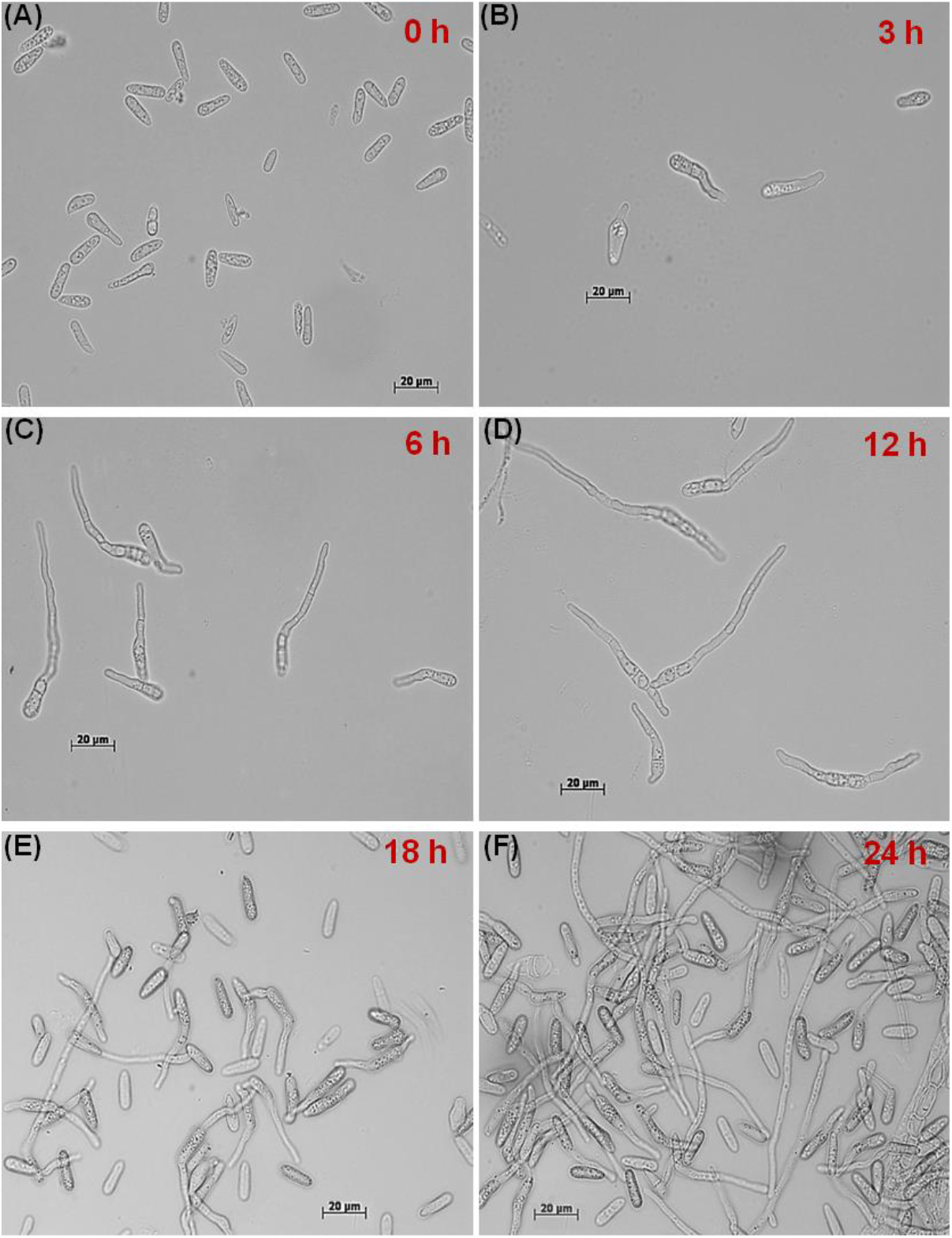
Different stages of GT formation at different time points in *C. gloeosporioides in-vitro*. **A)** Swelling and adhesion of conidia. **B)** GT initiation. **C), D) and E)** GT elongation. **F)** Hyphal network. Scale Bar = 20μm.

### 3.2. CAT fusion and GT formation are dependent on conidial density

A density of 4×10^4^ and 4×10^5^ conidia per ml of water was found to be optimal for CAT fusion in *C. gloeosporioides*. The CAT fusion percentages were 9.8% ± 3% and 12.3 ± 3.4% in 4×10^4^ and 4×10^5^ conidia per ml of water, respectively (Fig. 1D). However, for GT formation 4×10^5^ conidial density was optimal, resulted in 21.2% ± 1% GT formation (Fig. 1E). GT formation decreased to 8% ± 1% when the conidial density reduced to 10-fold that is 4×10^4^ conidia per ml of PDB (Fig. 1E). Very less and/or high conidial densities *viz*. 4×10^2^, 4×10^3^ and 4×10^6^ were not favorable for GT as well as CAT formation. These results suggest that CAT fusion and GT formation were dependent on conidial number threshold.

### 3.3. Differential nutritional requirements for CAT versus GT formation

The conidia of *C. gloeosporioides* failed to undergo CAT fusion in the presence of nutrient rich medium like PDB; and under heat and antifungal stress (Fig. 1F). As shown in the previous section, the high percentage of CAT fusion (13.4% ± 1.5%) was observed in sterile distilled water. However, when the conidia were incubated in water with 2% glucose, the CAT fusion frequency decreased to 3.4% ± 0.5% (Fig. 1F). It shows that CAT fusion occurred predominantly during nutrient starvation conditions and gets inhibited in the presence of nutrients. The conidia failed to induce CAT fusion in the presence of InSolution^™^ PD98059, an inhibitor of MEK, thereby suggesting the involvement of MAPK pathway in CAT fusion in *C. gloeosporioides* (Fig. 1F).

On the other hand, GT formation in *C. gloeosporioides* conidia was observed in presence of nutrients like PDB and 2% Glucose at the frequency of 20.3% ± 0.2% and 15% ± 1.4%, respectively (Fig. 1F). However, GT formation was not seen in nutrient limiting conditions like water as a medium, which indicates that GT formation occurs predominantly in nutrient rich environment but not in nutrient limiting conditions. Surprisingly, GT formation frequency increased up to 25% ± 1.3% in presence of the InSolution™ PD98059, an inhibitor of MEK (Fig. 1F).

### 3.4. RNA-Seq general data analysis

All RNA samples had an RNA integrity number (RIN) greater than 9.0. An average of 41.62 million raw sequencing PE reads were produced in total for three samples, out of which an average of 39 million reads were used for the downstream analysis after preprocessing. For every sample on an average of 95% of high-quality data was retained and used for analysis (Supplementary table 1). A total of 2,11,330 transcripts were obtained, which consisted of 73187, 64639 and 73504 transcripts of resting conidia, GT and CAT respectively. Among these transcripts, 35,373 were present in all three life stages, while 22522, 16930 and 24509 transcripts were uniquely expressed in resting conidia, GT and CAT, respectively (Fig. 4).

**Fig. 4:**
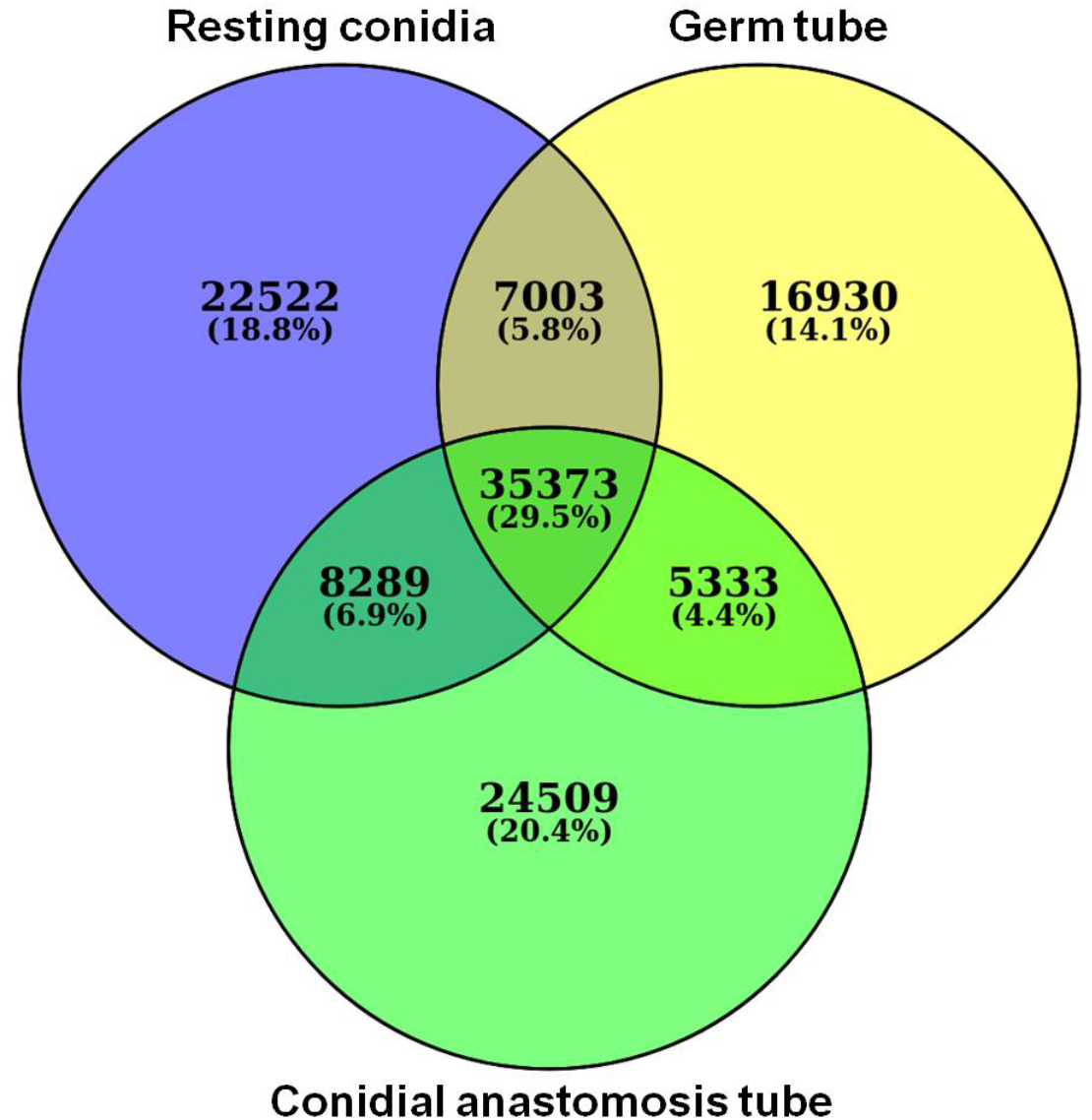
Venn diagram showing number of transcripts in resting conidia, GT and CAT.

The assembly of high-quality reads was performed separately for resting conidia, GT and CAT. Sample wise transcripts were merged and further clustered into 1,21,260 transcripts with an average length of 603 bp and N50 of 1849 bp (Supplementary table 2). Length of the final assembly was ranging from 37 to 31011 bps.

### 3.5. Gene ontology enrichment analysis of GT and CAT

The assembled transcripts for GT and CAT were individually grouped into ten molecular functions (MF) and ten biological processes (BP) (Fig. 5). The transcripts, which were significantly enriched and variable in numbers under the biological processes included fungal type (GO:0031505), carbohydrate (GO:0005975) and transmembrane (GO:0055085); and five molecular functions included. chitin (GO:0008061), serine type (GO:0004252) iron (GO:0005506), oxidoreductase (GO:0016491) and hydrolase (GO:0004553) during both GT formation and CAT fusion (Fig. 5). Biological process comparison of GT and CAT indicated that fungal type (GO:0031505) GO terms were higher in CAT compared to GT, whilst carbohydrate (GO:0005975) GO terms were greater in GT compared to CAT (Fig. 5A-B). However, when the molecular functions were compared, it was observed that the serine type (GO:0004252) and iron (GO:0005506) GO terms were higher in CAT than GT (Fig. 5C-D).

**Fig. 5:**
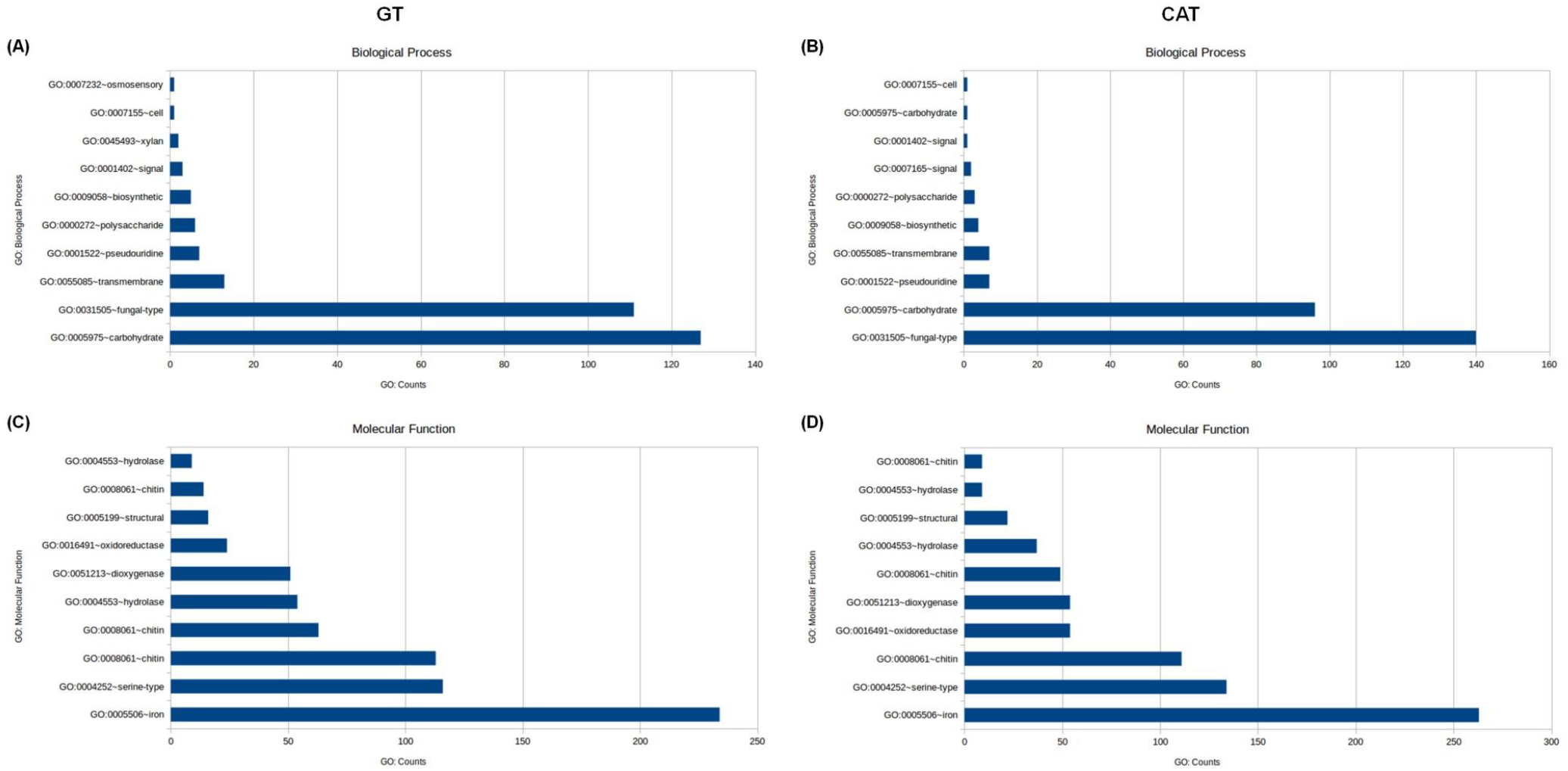
Gene ontology enrichment of biological processes and molecular functions during GT and CAT fusion in *C. gloeosporioides*. **A)** and **B)** GO counts of highly enriched GO biological processes terms in GT and CAT, respectively. **C)** and **D)** GO counts of highly enriched GO molecular function terms in GT and CAT, respectively.

### 3.6. Identification and functional annotation of DEGs in GT and CAT

The DEGs were further analyzed between GT and CAT. The transcripts which show a log2fold change less than −1 are represented as down regulated, the values greater than 1 are represented as up regulated and between −1 to 1 are termed as neutrally regulated.

Heatmaps of top 50 differentially expressed genes in GT and CAT showed a significant variation in their expression profiles. The top 25 up-regulated transcripts of GT showed significant down regulation in CAT. On the other hand, the top 25 up-regulated gene in CAT were found to be significantly down regulated in GT (Fig. 6). Heatmaps based on DEGs of resting conidia v/s GT and resting conidia v/s CAT also showed remarkable differential expression in these life stages of *C. gloeosporioides* (Supplementary Fig. 1). Manual functional annotation of top 50 transcripts for both GT and CAT resulted in certain categories of proteins, which probably play an important role in development of CAT and GTs in *C. gloeosporioides* (Supplementary table 3 and 4). Out of these top 50 differentially expressed transcripts, we found four major groups of highly up-regulated genes during GT formation, which were involved in cell wall degradation, host-fungus interaction, germination, transport, signaling and virulence, whose possible functions and expression values in GT and CAT are given in Table 1. Likewise, out of those top 50 DEGs, four major groups of highly up-regulated genes involved in CAT fusion were stress response, cell wall, membrane transport and signaling, cytoskeleton, cell cycle and cell rescue and pathogenicity genes, whose possible functions and expression values in CAT and GT are given in Table 2.

**Fig. 6:**
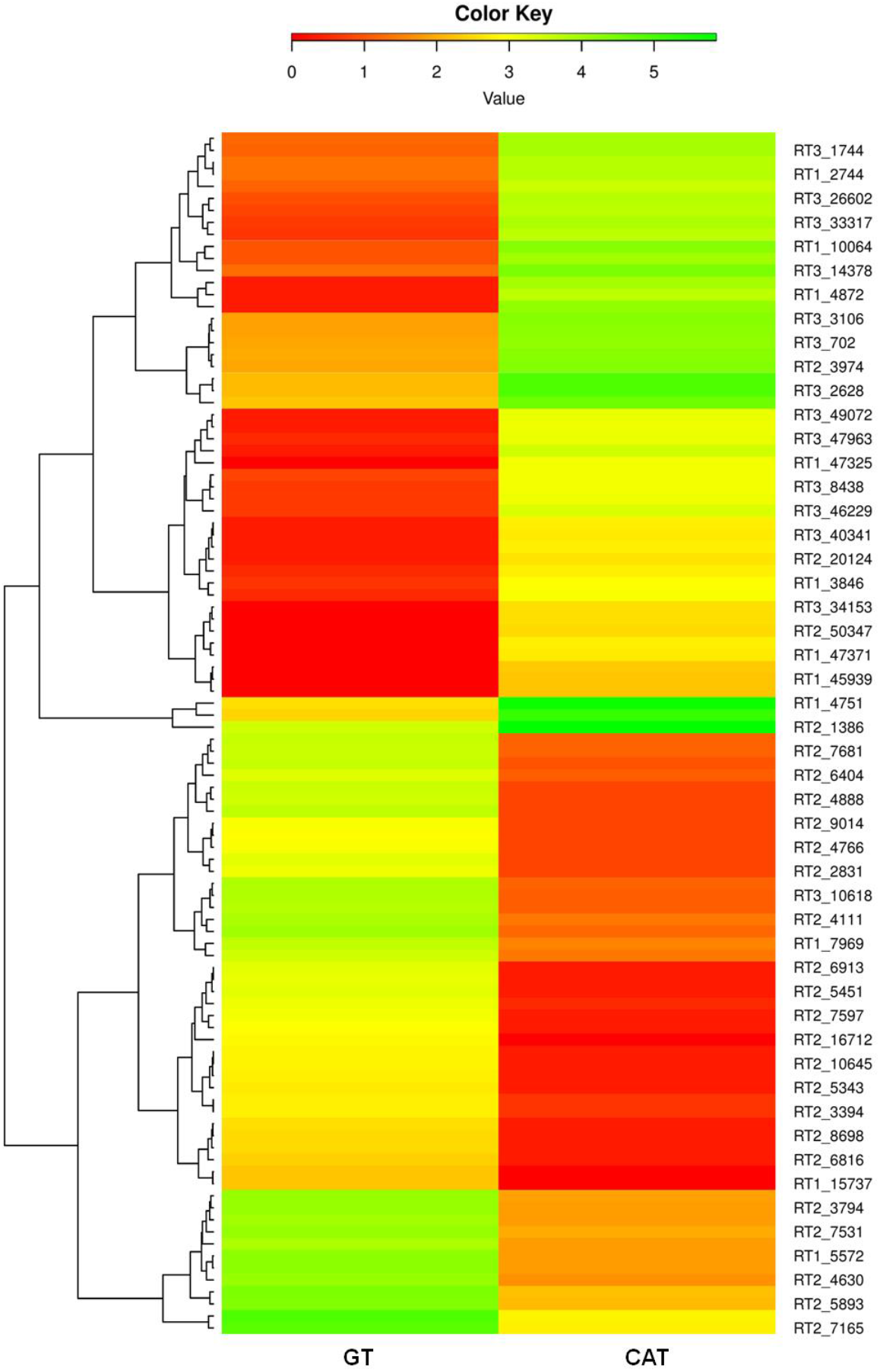
Heatmap depicting differential expression profile of selected genes involved during GT formation versus CAT fusion in *C. gloeosporioides*.

**Table 1:** Selected up-regulated genes during GT formation showing their possible functions and respective expression values in GT and CAT.

| Transcript ID | Gene name | Possible functions | GT | CAT |
| --- | --- | --- | --- | --- |
| **Cell wall degrading enzymes** |  |  |  |  |
| RT2_5451 | Pectin lyase | Degrades pectin | 1712 | 2 |
| RT2_4111 | Glycosyl hydrolase family 76 | Degrades cellulase, hemicellulose and lignin found in plant cell wall | 7920 | 24 |
| RT2_6404 | Mandelate racemase/muconate lactonizing enzyme domain-containing protein | Involved in the breakdown of lignin-derived aromatics | 1848 | 13 |
| RT2_6816 | Fungal cellulose binding domain-containing protein | Active in plant cell-wall hydrolysis | 254 | 2 |
| **Host-fungus interaction and germination** |  |  |  |  |
| RT2_16712 | Quinone reductase | Host-fungus interaction & infection | 761 | 1 |
| RT1_10305 | Hydrophobin 2 | Involved in adhesion | 9563 | 16 |
| RT1_4917 | Cerato-platanin | Role during fungus-plant interactions and infection | 16867 | 62 |
| RT2_5893 | GPI-anchored cell wall beta-1,3-endoglucanase EglC | Role in germination | 21332 | 143 |
| RT3_7789 | Hydrophobic surface binding protein A | Increase the hydrophobicity of conidia, aerial hyphae and fruiting bodies | 86745 | 621 |
| **Transport and signaling** |  |  |  |  |
| RT2_6913 | Fg-gap repeat protein | Important for ligand binding | 1571 | 2 |
| RT2_4144 | Phosphate-repressible phosphate permease/ transporter | Membrane transport proteins | 1556 | 2 |
| RT2_4587 | Transmembrane amino acid transporter | Ubiquitin dependent endocytosis and signaling responses | 3406 | 6 |
| RT1_15737 | MFS monosaccharide transporter | Membrane transport proteins | 182 | 1 |
| RT2_4766 | High affinity methionine permease | Cysteine and methionine transport across plasma membrane | 1027 | 6 |
| RT3_9669 | Uso1 / p115 like vesicle tethering protein | Intracellular protein transport | 24227 | 173 |
| **Virulence** |  |  |  |  |
| RT2_3663 | Peptidase S41 family protein | Involved in pathogenesis | **4430** | **6** |
| RT2_27038 | nrps-like protein | Involved in virulence | **1299** | **3** |
| RT2_3495 | Polysaccharide deacetylase | Involved in virulence | **3561** | **10** |
| RT2_10908 | FGGY-family pentulose kinase | Involved in virulence | **625** | **2** |
| RT1_5572 | Acetylornithine aminotransferase | Required for growth, conidiogenesis and pathogenicity | **17456** | **64** |
| RT2_5240 | GMC oxidoreductase | Role in pathogenicity | **324** | **2** |
| RT2_2131 | GNAT family acetyltransferase | Role in chitin metabolism of fungi, development and pathogenicity | **10433** | **68** |
| RT2_7531 | CAP-22 protein | Expressed in the conidium during appressorium formation | **13700** | **100** |
| RT1_7969 | NmrA family transcriptional regulator | Required for the invasive virulence | **4225** | **33** |
| RT1_7368 | Cyanide hydratase | Catalyzes the hydration of cyanide to formamide. May be important for plant pathogenic fungi in infection of cyanogenic plants | **2902** | **23** |

**Table 2:**
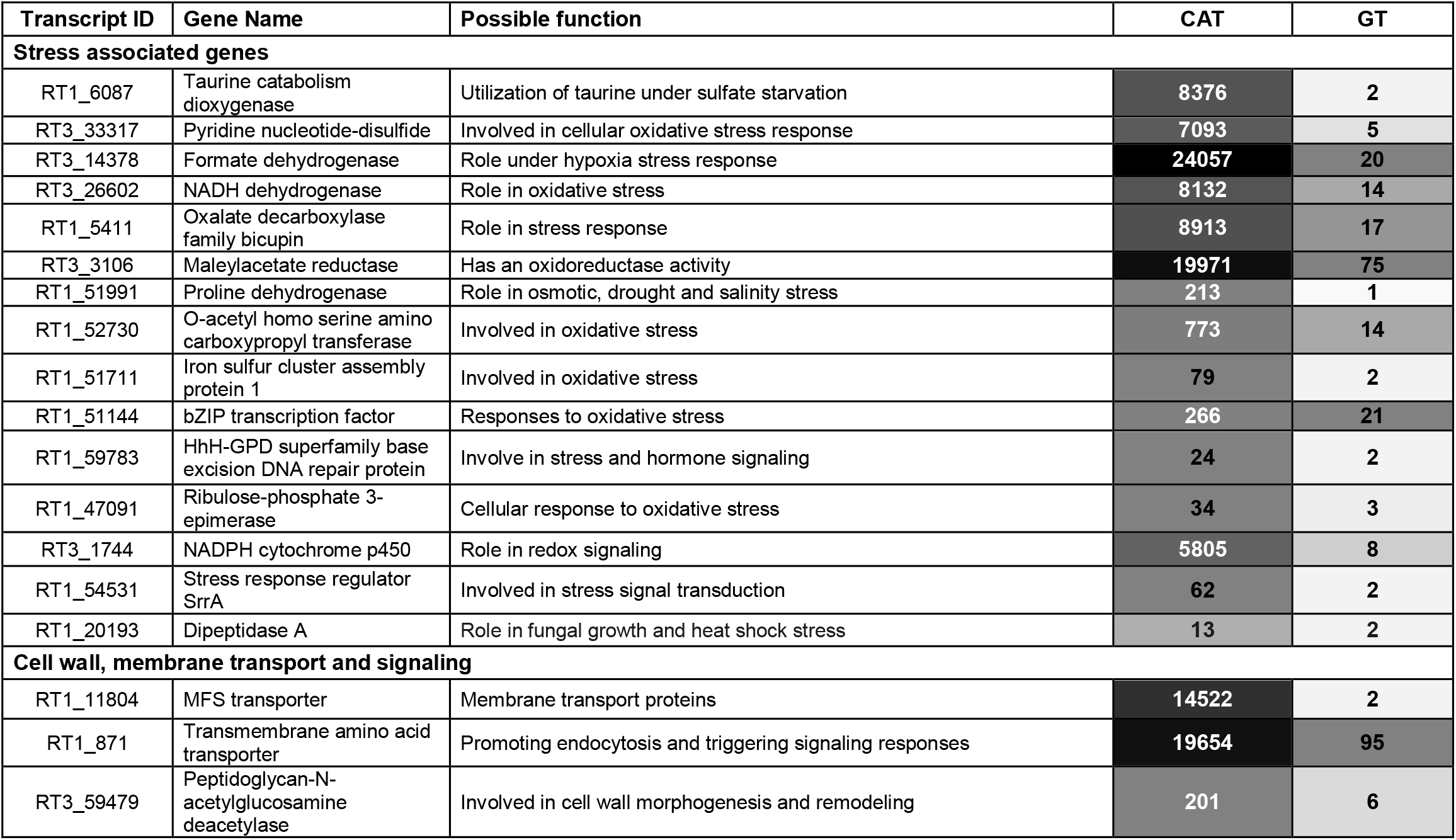
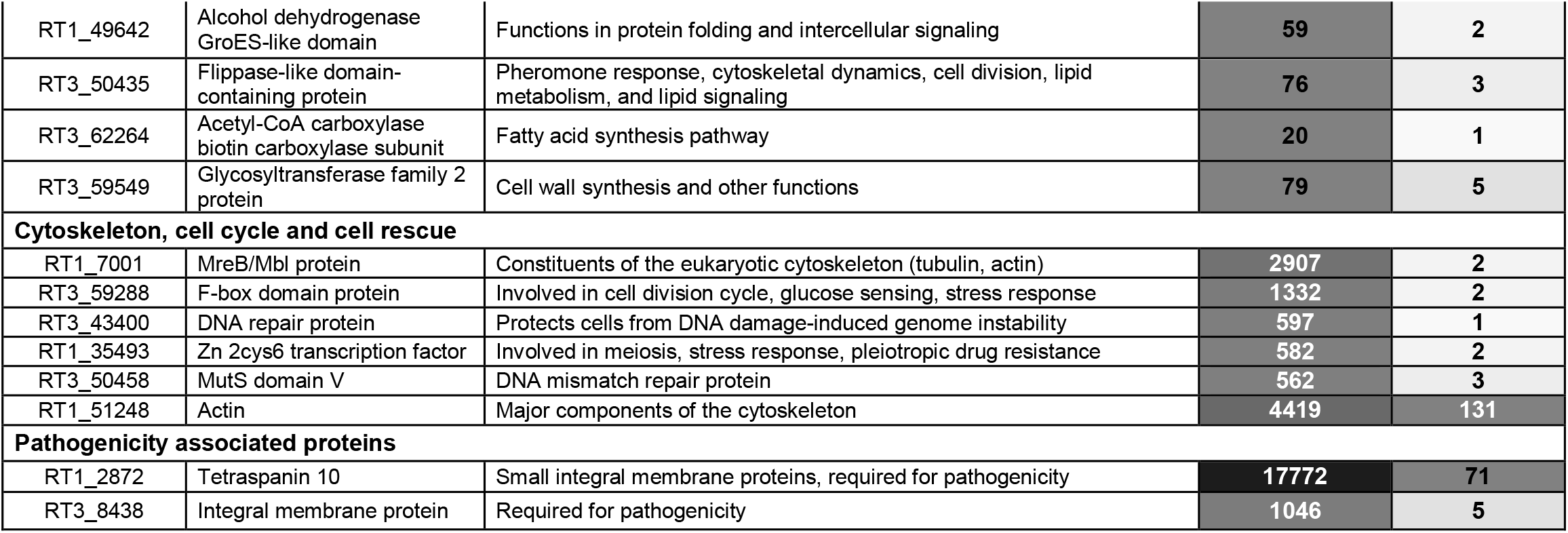
Selected up-regulated genes during CAT fusion, their possible functions and respective expression values in CAT and GT.

| Transcript ID | Gene Name | Possible function | CAT | GT |
| --- | --- | --- | --- | --- |
| **Stress associated genes** |  |  |  |  |
| RT1_6087 | Taurine catabolism dioxygenase | Utilization of taurine under sulfate starvation | 8376 | 2 |
| RT3_33317 | Pyridine nucleotide-disulfide | Involved in cellular oxidative stress response | 7093 | 5 |
| RT3_14378 | Formate dehydrogenase | Role under hypoxia stress response | 24057 | 20 |
| RT3_26602 | NADH dehydrogenase | Role in oxidative stress | 8132 | 14 |
| RT1_5411 | Oxalate decarboxylase family bicupin | Role in stress response | 8913 | 17 |
| RT3_3106 | Maleylacetate reductase | Has an oxidoreductase activity | 19971 | 75 |
| RT1_51991 | Proline dehydrogenase | Role in osmotic, drought and salinity stress | 213 | 1 |
| RT1_52730 | O-acetyl homo serine amino carboxypropyl transferase | Involved in oxidative stress | 773 | 14 |
| RT1_51711 | Iron sulfur cluster assembly protein 1 | Involved in oxidative stress | 79 | 2 |
| RT1_51144 | bZIP transcription factor | Responses to oxidative stress | 266 | 21 |
| RT1_59783 | HhH-GPD superfamily base excision DNA repair protein | Involve in stress and hormone signaling | 24 | 2 |
| RT1_47091 | Ribulose-phosphate 3-epimerase | Cellular response to oxidative stress | 34 | 3 |
| RT3_1744 | NADPH cytochrome p450 | Role in redox signaling | 5805 | 8 |
| RT1_54531 | Stress response regulator SrrA | Involved in stress signal transduction | 62 | 2 |
| RT1_20193 | Dipeptidase A | Role in fungal growth and heat shock stress | 13 | 2 |
| **Cell wall, membrane transport and signaling** |  |  |  |  |
| RT1_11804 | MFS transporter | Membrane transport proteins | 14522 | 2 |
| RT1_871 | Transmembrane amino acid transporter | Promoting endocytosis and triggering signaling responses | 19654 | 95 |
| RT3_59479 | Peptidoglycan-N-acetylglucosamine deacetylase | Involved in cell wall morphogenesis and remodeling | 201 | 6 |
| RT1_49642 | Alcohol dehydrogenase GroES-like domain | Functions in protein folding and intercellular signaling | 59 | 2 |
| RT3_50435 | Flippase-like domain-containing protein | Pheromone response, cytoskeletal dynamics, cell division, lipid metabolism, and lipid signaling | 76 | 3 |
| RT3_62264 | Acetyl-CoA carboxylase biotin carboxylase subunit | Fatty acid synthesis pathway | 20 | 1 |
| RT3_59549 | Glycosyltransferase family 2 protein | Cell wall synthesis and other functions | 79 | 5 |
| **Cytoskeleton, cell cycle and cell rescue** |  |  |  |  |
| RT1_7001 | MreB/Mbl protein | Constituents of the eukaryotic cytoskeleton (tubulin, actin) | 2907 | 2 |
| RT3_59288 | F-box domain protein | Involved in cell division cycle, glucose sensing, stress response | 1332 | 2 |
| RT3_43400 | DNA repair protein | Protects cells from DNA damage-induced genome instability | 597 | 1 |
| RT1_35493 | Zn 2cys6 transcription factor | Involved in meiosis, stress response, pleiotropic drug resistance | 582 | 2 |
| RT3_50458 | MutS domain V | DNA mismatch repair protein | 562 | 3 |
| RT1_51248 | Actin | Major components of the cytoskeleton | 4419 | 131 |
| **Pathogenicity associated proteins** |  |  |  |  |
| RT1_2872 | Tetraspanin 10 | Small integral membrane proteins, required for pathogenicity | 17772 | 71 |
| RT3_8438 | Integral membrane protein | Required for pathogenicity | 1046 | 5 |

### 3.7. Transcription factor candidates involved in GT formation and CAT fusion

In this study, a total of 49 Transcription factor (TF) families and 1044 TF proteins were predicted using the Fungal Transcription Factors Database; and out of them 38 TF families (Table 3) and 220 TF proteins (Supplementary table 5) were differentially expressed during GT and CAT induction. Twenty-four TF families e.g. bZIP, C2H2 zinc finger, HMG, Zn2Cys6 etc. were up-regulated during CAT fusion compared to GT formation, whilst 14 TF families such as AraC type, Homeobox etc. were up-regulated in GT formation (Table 3). Among 220 transcription factor genes, 42 and 82 were uniquely expressed during GT formation and CAT fusion, respectively (Supplementary table 5).

**Table 3:**
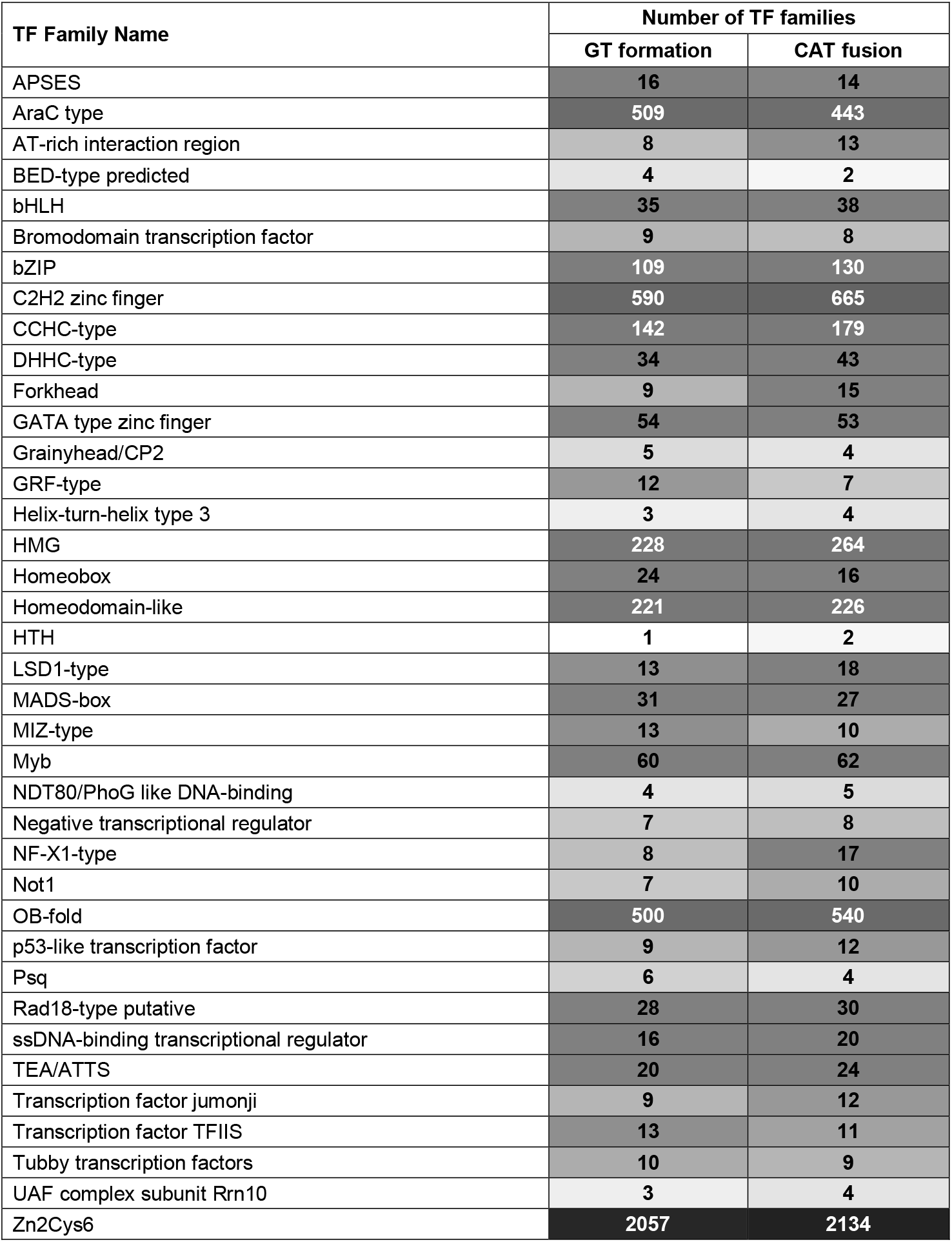
Transcription factors families and their numbers involved in GT formation and CAT fusion.

### 3.8. Effector candidates secreted during GT formation and CAT fusion

GT formation as well as CAT fusion in *C. gloeosporioides* involves deployment of secreted effector proteins. Effector candidates are often < 200 amino acids in length and cysteine-rich (Oliva *et al*., 2010), therefore, we principally focused on small secreted proteins < 250 amino acids in length. There was total 1567 effector candidates, of them 776 and 791 were secreted during GT and CAT formation, respectively. Out of 776 GT effector proteins, 103 were uniquely secreted during GT formation (Supplementary table 6) and out of 791 CAT effector proteins, 101 were uniquely secreted during CAT fusion (Supplementary table 7).

Differentially expressed secretory proteins were manually annotated and categorized in four and three major groups for GT formation and CAT fusion, respectively. Few selected uniquely expressed effector candidates during GT formation include hydrolytic enzymes, adhesion, germination, hyphal development associated proteins, transport, signaling related proteins, and proteins responsible for virulence. Some examples of these groups of effector proteins uniquely secreted during GT formation and their possible functions are given in Table 4. Likewise, the major effector proteins uniquely secreted during CAT fusion included stress associated proteins, cell wall, membrane transport, signaling related proteins, cytoskeleton, cell cycle, and fungal development associated proteins. Some examples of these groups of effector proteins uniquely secreted during CAT fusion and their possible functions are given in table 5.

**Table 4:** Uniquely expressed effector secretory proteins during GT formation.

| Transcript_ID | Protein names | Possible functions |
| --- | --- | --- |
| **Hydrolytic enzymes** |  |  |
| RT2_4955.p1 | Acetyl esterase | Degradation of hemicelluloses and pectin |
| RT2_12928.p2 | Alkaline phosphatase | Role in P mobilisation from organic substrates under P starvation conditions |
| RT2_9649.p1 | Amidohydrolase | Type of hydrolase that acts upon amide bonds |
| RT2_4824.p1 | Berberine bridge enzyme | Is a cellobiose oxidase which degrades the carboxymethylcellulose (CMC), xylan and lignin |
| RT2_14227.p1 | Beta-1,3-endoglucanase | Role in cell wall softening |
| RT2_10729.p1 | Endo-chitosanase | Role in degradation of the deacetylated portion of chitin in the fungal cell wall |
| RT2_9596.p1 | Extracellular cellulase | Degrades cellulose and some other related polysaccharides |
| RT2_1942.p1 | Glycosyl hydrolase family 15 | Degrades cellulase, hemicellulose and lignin found in plant cell wall |
| RT2_15512.p2 | Glycosyl hydrolase family 28 | Degrades cellulase, hemicellulose and lignin found in plant cell wall |
| RT2_618.p1 | Glycosyl hydrolase family 31 | Degrades cellulase, hemicellulose and lignin found in plant cell wall |
| RT2_12327.p1 | Xylosidase arabinofuranosidase | Role in xylan degradation |
| **Adhesion, germination and hyphal development** |  |  |
| RT2_1963.p1 | Acid trehalase | Provides energy during conidial germination |
| RT2_6726.p1 | Carbonic anhydrase | Role in fruiting body development and ascospore germination |
| RT2_5938.p1 | Developmentally Regulated MAPK Interacting protein | Role in diverse stresses and developmental processes |
| RT2_2892.p2 | Endoglucanase 3 | Role in germination |
| RT2_10852.p2 | Fungal hydrophobin | Role in adhesion |
| RT2_5434.p1 | Glycosyltransferase sugar-binding region containing DXD motif | Role in pathogenesis of plants by enabling hyphal growth |
| RT2_4975.p1 | Manganese lipooxygenase | Accelerates programmed spore germination |
| RT2_13809.p1 | Pollen proteins Ole e I like | Potentially implicated in pollen germination |
| RT2_18566.p1 | Spindle poison sensitivity protein | Role as spindle machinery for chromosome segregation and cytokinesis |
| RT2_10806.p1 | WD40 domain protein beta propeller | Regulates fungal cell differentiation |
| RT2_11269.p2 | Acid phosphatase, putative | Involved in fungal growth |
| RT2_4368.p1 | Class III aminotransferase, putative | Regulate the fungus-host interaction |
| **Transport and signaling** |  |  |
| RT2_12723.p2 | Bacterial extracellular solute-binding protein, family 3 | Act as chemoreceptors, recognition constituents of transport systems, and initiators of signal transduction pathways |
| RT2_18223.p1 | Eukaryotic porin | It is an important regulator of Ca<sup>2+</sup> transport in and out of the mitochondria |
| RT2_1739.p1 | WSC domain-containing protein | Responses to stress cues and metal ions |

| **Virulence** |  |  |
| --- | --- | --- |
| RT2_4851.p1 | Calcineurin-like phosphoesterase | Role in stress survival, sexual differentiation, and virulence |
| RT2_11269.p2 | Phosphoinositide phospholipase C, Ca<sup>2+</sup>-dependent | Roles in growth, stress tolerance, sexual development, and virulence |
| RT2_3804.p1 | Secretory lipase, putative | Potential virulence factors |
| RT2_5435.p1 | Versicolorin b synthase | Role in biosynthesis of aflatoxins |
| RT2_15735.p1 | Zinc-binding dehydrogenase | Role in infection |

**Table 5:** Unique secretory effector candidate genes expressed during CAT fusion.

| Transcript_ID | Gene names | Possible functions |
| --- | --- | --- |
| **Stress associated** |  |  |
| RT3_6147_m.31990 | Acetyl-CoA acetyltransferase | Role in oxidative and cell wall stresses |
| RT3_8826_m.37297 | Calcineurin-like phosphoesterase | Involved in stress survival and fungal virulence |
| RT3_10144_m.39239 | Chitin synthesis regulation, resistance to Congo red | Role in cell wall stress, noxious chemicals and osmotic pressure changes |
| RT3_12971_m.42487 | Cytochrome oxidase assembly protein | Cox11p is an integral protein of the inner mitochondrial membrane that is essential for cytochrome c oxidase assembly. Role in response to hydrogen peroxide exposure |
| RT3_12332_m.41840 | LipA and NB-ARC domain-containing protein | Function as key integrators of stress and nutrient availability signals |
| RT3_571_m.6736 | Pyridoxamine 5'-phosphate oxidase | Role in oxidative stress |
| RT3_15296_m.44520 | Spherulation-specific family 4 | Required for spherulation under starvation conditions |
| RT3_2386_m.18522 | SWIM zinc finger | Roles in the stress response and virulence |
| RT3_10404_m.39608 | WSC domain-containing protein | Responses to stress cues and metal ions |
| **Cell wall, membrane transport and signaling** |  |  |
| RT3_17141_m.45950 | Ankyrin-3-like protein 3 | Roles in cell motility, activation, proliferation, contact, and the maintenance of specialized membrane domains |
| RT3_870_m.9203 | Aspartic endopeptidase | Role in cell-wall assembly, remodeling and cell wall integrity |
| RT3_2825_m.20686 | Blastomyces yeast-phase-specific protein | Required for cell wall morphogenesis and pathogenesis |
| RT3_100_m.1747 | Cytochrome p450 | Role in redox signaling |
| RT3_6853_m.33623 | Endo-beta-1,6-glucanase | Cell wall morphogenesis and remodeling |
| RT3_3953_m.25310 | Gamma-glutamyltranspeptidase | Role in the vacuolar transport and metabolism of glutathione |
| RT3_2987_m.21396 | GPI biosynthesis protein family Pig-F | Required for Cell Wall Biogenesis and Normal Hyphal Growth |
| RT3_25_m.564 | Hkr1p | Regulate the filamentation and mitogen-activated protein kinase pathway |
| RT3_2892_m.20978 | Mid2 like cell wall stress sensor | Required for stress-induced nuclear to cytoplasmic translocation of Cyclin C and cell wall integrity signaling pathway |
| RT3_18063_m.46545 | MIP family channel protein | Involved in the transport of water and neutral solutes across the membranes |
| RT3_268_m.3768 | NCS1 nucleoside transporter | Responsible for uptake of purines, pyrimidines or related metabolites |
| RT3_704_m.7811 | PAN domain | Essential for RasA-mediated morphogenetic signaling |
| RT3_5718_m.30849 | Pectin esterase | Facilitate plant cell wall modification and subsequent breakdown |
| RT3_18612_m.46914 | PQ loop repeat | Function as cargo receptors in vesicle transport |
| RT3_11944_m.41457 | Probable lipid transfer | Involved in lipid transfer |
| RT3_21269_m.48652 | Protein serine/threonine kinase | Roles in cellular function like regulation of signaling pathways |
| RT3_3541_m.23752 | Protein YOP1 | Involved in membrane/vesicle trafficking |
| RT3_543_m.6478 | Proteolipid membrane potential modulator | Pmp3 is transcriptionally regulated by the Spc1 stress MAPK (mitogen-activated protein kinases) pathway |
| RT3_2326_m.18214 | SOCE-associated regulatory factor of calcium homeostasis | Negative regulator of intracellular calcium signaling |
| RT3_2348_m.18321 | Stretch-activated Ca<sup>2+</sup>-permeable channel component | Required for Ca<sup>2+</sup> influx induced by the mating pheromone, alpha-factor |
| RT3_13567_m.43062 | Tetraspanin | Superfamily of small integral membrane proteins and required for pathogenicity |
| **Cytoskeleton, cell cycle and fungal development** |  |  |
| RT3_12912_m.42432 | Alkaline proteinase | Role in fungal physiology and development |
| RT3_20705_m.48270 | Coronin | Role in the organization and dynamics of actin, F-actin remodeling |
| RT3_13797_m.43265 | Frequency clock protein | Regulates various aspects of the circadian clock in *Neurospora crassa* |
| RT3_16749_m.45643 | Galactoside-binding lectin | Role in fruiting body development |
| RT3_15066_m.44317 | Nucleoside diphosphate kinase | Regulation of spore and sclerotia development |
| RT3_1_m.4 | WD40 domain-containing protein | Regulates fungal cell differentiation |
| RT3_3897_m.25109 | Wlm domain-containing protein (Fragment) | Related to the DNA damage response proteins WSS1 involved in sister chromatid separation and segregation |

### 3.9. Real time qPCR validation of selected DEGs

Differentially expressed genes in resting conidia, GT and CAT identified by RNA-seq were further verified through a quantitative real time PCR (qRT-PCR). Log2fold expression values were used to evaluate correlation between RNA-seq and qRT-PCR. The results obtained by qRT-PCR showed significant correlation (R^2^ = 0.736; *P*< 0.01) (Fig. 7). It showed the RNA-seq experiment was conducted with accuracy and RNA-seq data is reliable to further used to understand molecular mechanism underlying CAT formation. We observed a strong correlation between the qRT-PCR and RNA-Seq analysis of 17 DEGs in GT versus CAT formation (Fig. 8). Similar levels of correlations were also attained in expression levels of 14 and 13 DEGs in resting conidia v/s GT and resting conidia v/s CAT, respectively (Supplementary Fig. 2).

**Fig. 7:**
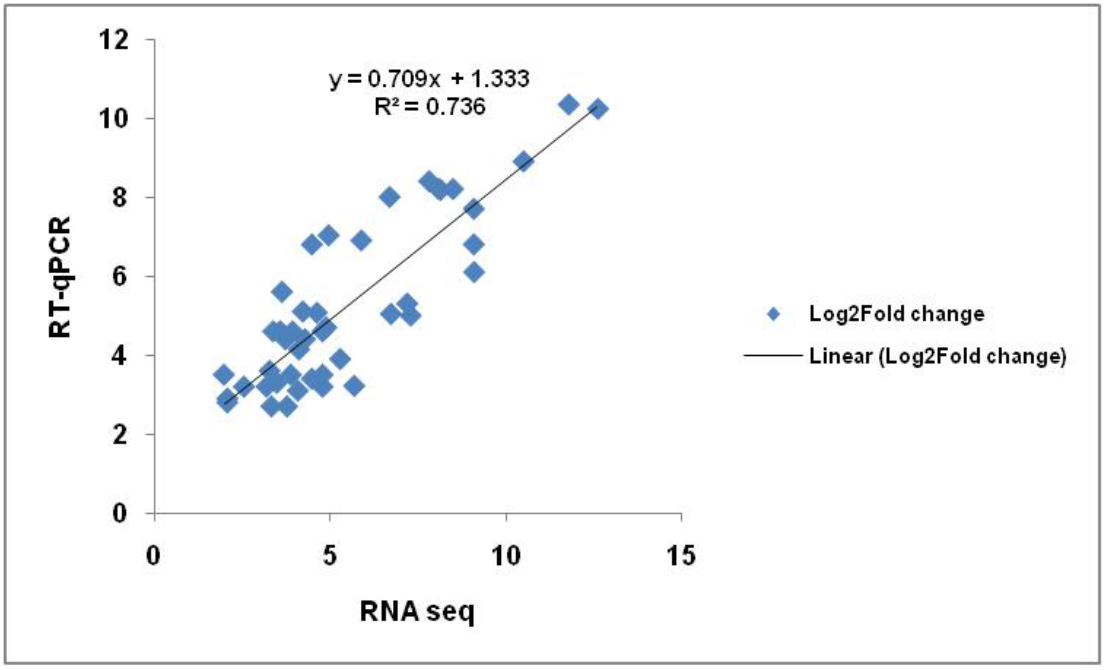
Scatter plot showing correlation between of expression levels of DEGs detected by qRT-PCR and RNA seq in GT and CAT in *C. gloeosporioides*.

**Fig. 8:**
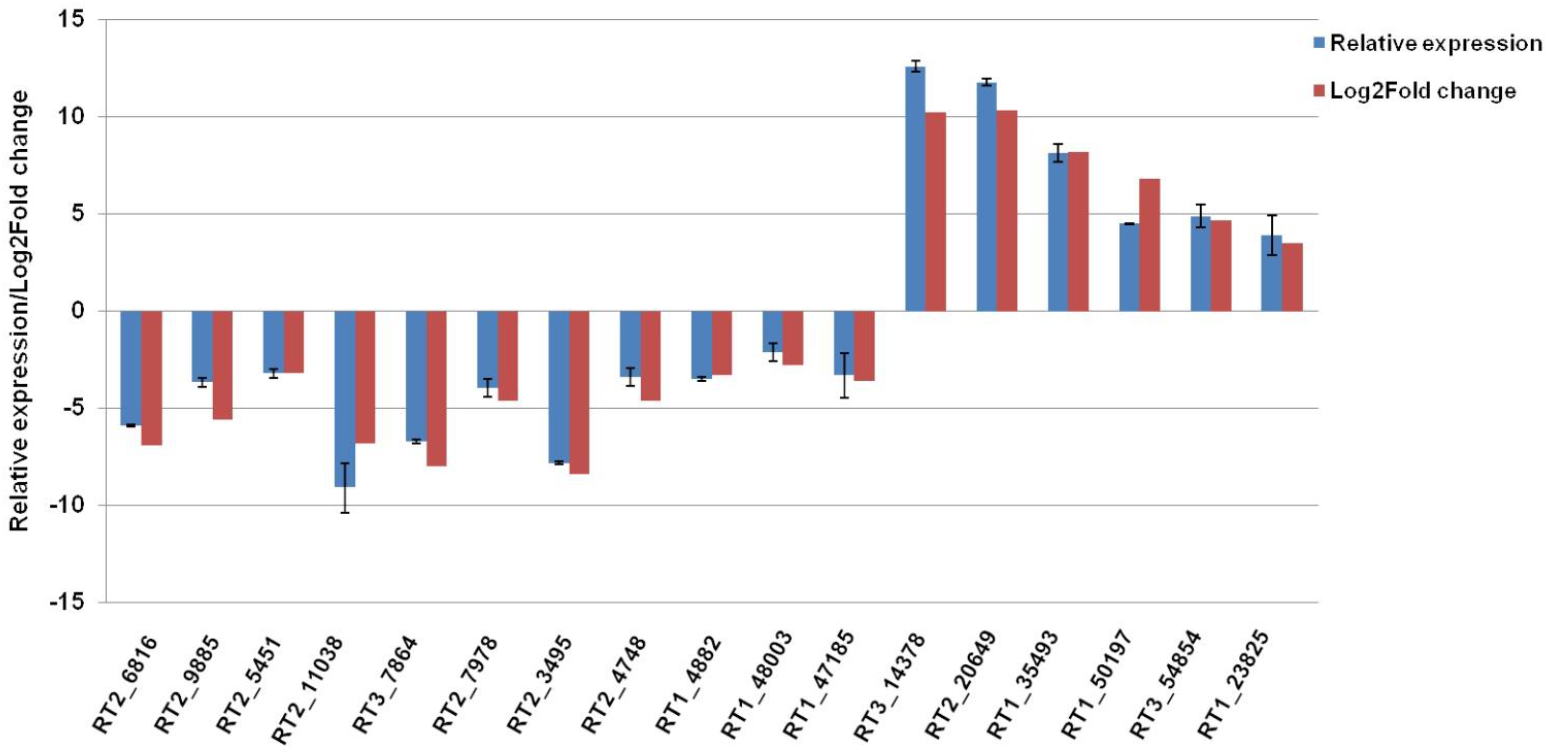
Comparison of differential expression of DEGs obtained by qRT-PCR and RNA-Seq in GT formation versus CAT fusion in *C. gloeosporioides*.

## 4. Discussion

### 4.1. Differential physiological requirements for GT and CAT

We have previously optimized the *in-vitro* conditions for an inter-specific CAT fusion between *C. gloeosporioides* and *C. siamense* (Mehta and Baghela, 2020). In present study, we have optimized the *in-vitro* conditions for GT and CAT induction in *C. gloeosporioides*. We observed significant CAT fusion in 17 d old conidia; and GT formation in 6 d old conidia (Fig. 1A). It has been reported that CAT fusion is dependent on conidial age for different species; 16 days old conidia of *C. lindemuthianum* and *C. gossypii* were more appropriate for CAT fusion, whilst 20 days old conidia of *C. fructicola* and *C. nymphaeae* were suitable for CAT fusion (Gonçalves *et al*., 2016; Roca *et al*., 2004, 2003), while the younger conidia (10 d old) of *C. lindemuthianum* were found to be suitable for GT formation in nutrient rich medium like PDB (Ishikawa *et al*., 2010). Howbeit, the conidial age requirements for both CAT fusion and GT formation in *F. oxysporum* and *N. crassa* were found to be similar 7-10 and 5-6 days, respectively (Kurian *et al*., 2018; Roca *et al*., 2005b; Shahi *et al*., 2016). Our result seems to fit with the conidial age requirement of the genus *Colletotrichum*; older conidia (17 d old) for CAT fusion and younger conidia (6 d old) for GT formation were preferred in *C. gloeosporioides* (Fig. 1A). The GT/CAT formation was shown to be conidial density dependent in *N. crassa* and *F. oxysporum* (Kurian *et al*., 2018; Roca *et al*., 2005a). In these fungi, the conidial number threshold requirement for CAT induction was more or less same i.e. 1×10^6^ conidia/ml (Gonçalves *et al*., 2016; Kurian *et al*., 2018; Leu L.S., 1967; Roca *et al*., 2005a, 2003; Shahi *et al*., 2016). The GT induction has also been shown to be dependent on conidial density in *F. oxysporum*, the maximum GT formation was observed in 1×10^5^ conidia per ml (Kurian *et al*., 2018). The conidial number requirement for CAT fusion and GT formation in *C. gloeosporioides* was also found to be 4×10^5^ conidia/ml (Fig. 1D-E). It indicates that a conidial threshold is a requisite for the CAT induction as well as GT formation (Fig. 1D-E).

It has been shown previously that nutrient limiting conditions induce CAT fusion and nutrient rich conditions promote GT formation in few fungi (Ishikawa *et al*., 2010; Kurian *et al*., 2018). We observed that the conidia of *C. gloeosporioides* could efficiently form CAT in the absence of nutrients (water) compared to nutrient rich conditions like PDB. Presence of glucose in water also reduces CAT fusion percentage in this fungus (Fig. 1F). On the other hand, significant GT formation occurred only in the presence of nutrients like PDB and nutrient limiting conditions negatively affects GT formation (Fig. 1F). These observations indicate that the CAT fusion occurs in nutrient limiting conditions; conversely GT formation occurs only in the nutrient rich conditions in *C. gloeosporioides*. In another species of the genus *Colletotrichum* i.e., *C. lindemuthianum*, CAT fusion doesn’t occur in PDB and even in nutrient poor Vogel’s medium, while it occurs only in water (Ishikawa et al. 2010). While in case of other CAT forming fungi like *N. crassa* and *F. oxysporum*, CAT fusion does not occur in water, while it requires some amount of nutrients for CAT fusion e.g., Vogel’s medium for *N. crassa* and YNB+KNO3 /1% PDB +NaNO3 for *F. oxysporum* (Fischer-Harman *et al*., 2012; Kurian *et al*., 2018; Palma-Guerrero *et al*., 2009; Roca *et al*., 2005a; Shahi *et al*., 2016). We have also shown that CAT fusion in *C. gloeosporioides* gets inhibited by a MAP kinase inhibitor InSolution^™^ PD98059 (Fig. 1F), thereby suggesting a possible role of MAP kinase pathway in CAT fusion in *C. gloeosporioides*. Interestingly, in the presence of MAP kinase pathway inhibitor InSolution™ PD98059, the frequency of GT formation increased considerably compared to PDB (Fig. 1F), thereby suggesting that this pathway may not be a significant player in early GT formation. However, the very same pathway was shown to be essential in appressorium formation in the same fungus (Wang *et al*., 2018).

We propose that the choice of conidia to either form GT or CAT depends upon multiple factors *viz*. conidial age (internal nutrients), availability of external nutrients and/or starvation stress conditions. We have shown that the older conidia were superlative for CAT fusion while younger conidia were ideal for GT formation. We believe that increased CAT fusion reduces the internal resources for GT formation; however, the conidia’s decision to either form CAT or GT do not only rely solely upon the internal resources. We have shown that when the older conidia (17 days) were incubated in water (no nutrients), they could undergo extensive CAT fusion but no GT formation, however when the same aged older conidia were incubated in PDB, they could also form GT but at a reduced rate compared to younger conidia (6 and 10 days old) (Fig. 1A). Therefore, we hypothesize that as the age of conidia increases, the internal resources get depleted, in this scenario, if the external resources are provided then conidia could still form GTs to a lesser extent. On the contrary, in the absence of external nutritional resources, the older conidia undergo extensive CAT fusion.

After understanding the differential physiological requirements of CAT and GT formation, we attempted to decipher the differential molecular requirements for these two processes in *C. gloeosporioides* by undertaking RNA-sequencing and analysis. We observed significant differences in transcriptomes of GT formation and CAT fusion. Further, we identified many differentially expressed genes, transcription factors and secretory effector proteins during GT formation and CAT fusion in *C. gloeosporioides*.

### 4.2. Transcriptome analysis of GT

Genes involved in cell wall degradation, host-fungus interaction, germination, transport, signalling, and virulence were found to be up-regulated during GT formation. Previously, it has been shown that gene responsible for plant cell wall degradation; secondary metabolism and detoxification were up-regulated during GT formation in *C. fructicola* (Zhang *et al*., 2018). We also observed that some hydrolytic enzymes involved in plant cell wall degradation such as pectin lyase, glycosyl hydrolase families, mandelate racemase/ muconate lactonizing enzyme and fungal cellulose binding domain-containing protein were up-regulated during GT formation (Table 1). Fungal hydrophobins are small, secreted hydrophobic proteins involved in the adhesion of conidia to the host surface prior to GT formation (Tucker and Talbot, 2001). Fungal hydrophobin, HsbA like protein were shown to be up-regulated during GT and appressorium development in *A. oryzae* and *C. fructicola* (Ohtaki *et al*., 2006; Zhang *et al*., 2018). In the present study, Hydrophobin 2 and HsbA genes were also up-regulated during GT formation in *C. gloeosporioides*, which probably be involved in conidial adhesion. Other significantly differentially expressed genes include GPI-anchored cell wall beta-1,3-endoglucanase Egl, which probably plays a role in germination and hydrophobic surface binding protein A, which increases the hydrophobicity of conidia, aerial hyphae and fruiting bodies.

Fungal transporter proteins play a broad range of biological functions, and an important role in pathogenicity of phytopathogens (Gupta and Chattoo, 2008; Stefanato *et al*., 2009; Sun *et al*., 2006). In previous studies, it has been demonstrated that an ABC protein CgABCF2 was required for appressorium formation and plant infection in *C. gloeosporioides* (Wang *et al*., 2018; Zhou *et al*., 2017). In the present study, few transporters and signalling genes *viz*. Fg-gap repeat protein, phosphate-repressible phosphate transporter, transmembrane amino acid transporter, MFS monosaccharide transporter, high affinity methionine permease and Uso1 / p115 like vesicle tethering protein were up-regulated during GT formation. These transports and associated signalling genes are probably important to sense the favourable conditions for germination, which results in GT formation.

Some secondary metabolites and virulence associated genes were shown to add to pathogenicity in *C. gloeosporioides* (Gan *et al*., 2013). We have also found out various secondary metabolite backbone genes like peptidase S41 family protein, nrps-like protein, FAD dependent oxidoreductase and cyanide hydratase were up-regulated during GT formation, which may contribute to pathogenicity (Le Govic *et al*., 2019; Muszewska *et al*., 2017). During GT formation, few other highly up-regulated genes such as polysaccharide deacetylase, FGGY-family pentulose kinase, acetylornithine aminotransferase, GMC oxidoreductase, GNAT family acetyltransferase, CAP-22 protein and NmrA family transcriptional regulator may also be implicated for pathogenicity in *C. gloeosporioides* (Table 1).

Transcription factors play important roles in various biological processes. Out of 220 TF genes, 42 were uniquely expressed during GT formation, and these genes are probably involved in regulation of GT formation in *C. gloeosporioides* (Supplementary table 5). Effector proteins are usually secretory proteins, which can play an important role in plant infection. Previous study has shown that effectors are secreted from appressorium before host invasion in *C. higginsianum*, a close relative of *C. gloeosporioides* (Kleemann *et al*., 2012). In our study, we have identified uniquely expressed effector candidate genes during GT formation. Interestingly, the annotated effectors could also be categorised in the same categories, which were based on DGE analysis of GT formation. The uniquely expressed effector gene candidates include hydrolytic enzymes, adhesion, germination, hyphal development associated proteins, transport, signalling related proteins, and proteins responsible for virulence (Table 4). Transcripts of gene glucanases that degrade glucan in the fungal cell wall for cell wall modulation were observed during germination in *Aspergillus niger* (van Leeuwen *et al*., 2013). Various transcripts and secretory effectors of glucanases were observed in our study during GT formation, which might be involved in the cell wall morphogenesis leading to germination. Trehalose is known to get accumulated in dormant conidia of *A. niger* and is degraded during germination (van Leeuwen *et al*., 2013). We detected trehalase as a secretory effector during GT formation, which probably provides energy by degrading trehalose in *C. gloeosporioides*. Other noteworthy uniquely expressed effector gene candidates include carbonic anhydrase and Manganese lipoxygenase, which might be important for fruiting body development, ascospore germination and accelerating programmed spore germination respectively (Table 4).

The evidences for involvement of MAPK in conidial germination are conflicting and equally diverse between the species. It is reported that deletion of MAPK gene CMK1 in *C. lagenarium* prevents germination, whereas deletion of MAPK gene pmk1 in *Magnaporthe grisea*, blocks appressorium development but not germination (Osherov and May, 2001). The MAPK pathway was shown to be essential for appressorium development in *C. gloeosporioides* (Wang *et al*., 2018). However, in our case, GT formation was not inhibited in the presence of MAPK pathway inhibitor Insolution™ PD98059, and moreover, no transcripts related to MAPK pathway could be detected in GT transcriptome, thereby indicating that MAPK pathway may not be essential for early GT formation in *C. gloeosporioides*.

### 4.3. Transcriptome analyses of CAT

The CAT fusion seems to involve a set of different molecular requirements compared to GT formation, while retaining few common transcripts in both of these processes. We observed that the major groups of highly up-regulated genes involved in CAT fusion belong to stress response, cell wall integrity, membrane transport, signalling, cytoskeleton, cell cycle and cell rescue processes (Table 2).

In this study, we found many stress associated genes were up-regulated during CAT fusion, e.g. pyridine nucleotide-disulfide, formate dehydrogenase, NADH dehydrogenase, oxalate decarboxylase family bicupin, maleylacetate reductase, O-acetyl homo serine amino carboxypropyl transferase, iron sulfur cluster assembly protein 1, bZIP transcription factor, HhH-GPD superfamily base excision DNA repair protein, ribulose-phosphate 3-epimerase and dipeptidase A. All these genes are believed to be involved in oxidative stress response. Reactive oxygen species (ROS) are known to play an important role in redox signalling pathways including cell differentiation, development, and cytoskeleton remodelling (Fischer and Glass, 2019). NADPH-oxidases (NOX) are responsible for production of superoxide (ROS) by oxidizing NADPH and reducing molecular oxygen, which is believed to be involved during communication and cell fusion in several fungi (Fischer and Glass, 2019). It has been shown that NADPH oxidase BcNoxA or Nox regulator BcNoxR gene deletion mutant of Grey mould *Botrytis cinerea* were defective in CAT fusion (Roca *et al*., 2012). In our study, NADPH oxidase cytochrome p450, and stress response regulator SrrA which probably participate in redox signalling and stress signal transduction were also found to be up-regulated during CAT fusion. It has been previously documented that starvation response is correlated with expression of genes encoding oxidative stress response in *S. cerevisiae* (Petti *et al*., 2011). Further oxidative stress was shown during aging of stationary cultures of *S. cerevisiae* (Jakubowski *et al*., 2000). In our study we have demonstrated that the older conidia under nutritional starvation exhibit CAT fusion and our transcriptome data analysis also revealed high expression of oxidative stress relates genes. This suggests that starvation in *C. gloeosporioides* conidia induced a strong oxidative response and the genes involved in this response might play important roles in CAT fusion in this fungus.

The CAT fusion involves sensing of quorum through signalling and transport followed by cell wall and cell membrane remodelling. Many membrane transporters and cell wall modulation genes including MFS transporter, transmembrane amino acid transporter, peptidoglycan-N-acetylglucosamine deacetylase, and Glycosyltransferase family 2 protein were found to be up-regulated during CAT fusion. Other important transcripts *viz*. alcohol dehydrogenase GroES-like domain, flippase-like domain-containing protein, and acetyl-CoA carboxylase were up-regulated during CAT fusion, which probably play roles in protein folding, pheromone response, cytoskeletal dynamics, cell division, lipid metabolism and fatty acid synthesis in *C. gloeosporioides*.

The cytoskeleton actin is important for several functions including cell polarity, exocytosis, endocytosis, septation, movement of organelle, and chemotropic growth of fungi (Fischer and Glass, 2019). In *N. crassa*, the actin cables and patches are significantly increased at CAT tips during chemotropic interactions between germlings or conidia (Roca *et al*., 2010). However, microtubules are dispensable for germling communication, while actin is critically important in *N. crassa* (Roca *et al*., 2010). We observed the MreB/Mbl proteins which are constituents of the eukaryotic actin cytoskeleton were highly up regulated during CAT fusion in *C. gloeosporioides*, suggesting that actin is critically important for polarized growth and CAT fusion. We have previously shown that an inter-specific CAT fusion between *C. gloeosporioides* and *C. siamense* involved nuclear transfer (Mehta and Baghela, 2020). Therefore, we assume that even in the present study, CAT fusion in *C. gloeosporioides* also involves nuclear exchange. Corroborating to this belief, we have observed that several cell cycle and cell rescue genes were highly up-regulated during CAT fusion in *C. gloeosporioides*, which include F-box domain gene, which is thought to be involved in mitosis or cell division cycle, DNA repair and MutS domain V gene, which is probably involved in protection of cells from DNA damage-induced genome instability by repairing the DNA mismatch.

Transcription factors play important roles in various biological processes. Out of 220 TF genes, 82 were uniquely expressed during CAT fusion, and these genes are probably involved in regulation of CAT fusion in *C. gloeosporioides* (Supplementary table 5). Two such transcription factors, adv-1 and pp-1, a C2H2-Zn2C transcription factor were previously shown to be necessary for germling communication and fusion in *N. crassa* (Dekhang *et al*., 2017; Leeder *et al*., 2013). Similarly, we have also observed that few transcription factors including C2H2-Zn2C, Zn 2cys6, bZIP transcription factor were up-regulated during CAT fusion in *C. gloeosporioides*.

We have also found various unique effector genes specifically secreted during CAT fusion and interestingly the major transcripts could be grouped in the similar categories like general DEGs, which included stress associated proteins, cell wall, membrane transport, signaling related proteins, cytoskeleton, cell cycle, and fungal development associated proteins. We observed few stress associated effector genes including Acetyl-CoA acetyl transferase, Calcineurin-like phosphoesterase, LipA and NB-ARC domain-containing protein and WSC domain-containing protein coding gene, which might be involved in physiological stress response and nutrient availability signals during CAT fusion in *C. gloeosporioides*. WSC domain-containing protein coding gene has previously been shown to be important in germling and hyphal fusion in *N. crassa* (Maddi *et al*., 2012). We observed few MAPK pathway related transcripts as secretory effectors during CAT fusion, which included Hkr1p that is known to regulate the filamentation and MAPK pathway, and PAN domain which is known to be essential for RasA-mediated morphogenetic signalling, and Pmp3 which is also involved in MAPK pathway. Up-regulation of MAP kinase pathway genes corroborates well with our physiological data on inhibition of CAT fusion in the presence of InSolution™ PD 98059, an inhibitor of MEK. These data together indicate that MAPK pathway is involved in CAT fusion in *C. gloeosporioides*. Calcium and some calcium dependent genes have shown to play a critical role during the membrane fusion in germling communication in various fungi (Fischer and Glass, 2019). Calcium is also essential for polarized hyphal growth and fusion in *N. crassa* (Palma-Guerrero *et al*., 2013; Takeshita *et al*., 2017). We detected some calcium dependent genes as secretory effectors, which included calcium influx-promoting protein ehs1, calcium-related spray gene, SOCE-associated regulatory factor of calcium homoeostasis gene, which are probably involved in maintaining cell wall integrity, calcium signalling, calcium stress response, growth and virulence during CAT fusion in *C. gloeosporioides*. We also detected some transcripts of cell division cycle genes, which uniquely secreted during CAT fusion, viz. coronin, which known to play role in the organization and dynamics of actin and F-actin remodelling, and Wlm domain-containing genes, which is associated with sister chromatid separation and segregation during mitosis.

## 5. Conclusion

We propose a model (created with BioRender.com) to explain the mutual exclusiveness of GT formation and CAT fusion and their dependency on conidial age, availability of external nutrients and differential RNA profiles (Fig. 9). As per the model, the younger conidia (6 days) are expected to have substantial internal resources, therefore, when such young conidia are incubated in a nutrient rich medium, they tend to form GT extensively, and when the same young conidia are incubated in a nutrient poor medium, they do not form GT and negligible CAT. On the other hand, when the older conidia (17 days) are believed to be exhausted of internal nutrients and therefore, when incubated in nutrient rich medium, they could form GT with very less frequency, however, when such older conidia are incubated in water (no nutrients), they tend to form extensive CATs (Fig. 9). During the GT formation genes responsible for adhesion, GT elongation, germination, were found to be highly up-regulated, which signifies that under suitable conditions, when the host plant is available for the conidia, they tend to undergo GT formation followed by infectious hyphae. The transcriptome data also revealed high expression of genes coding for hydrolytic enzymes, infectious hyphae and virulence factors during GT formation, which again signifies that eventually conidia will have to degrade the plant cell wall and establish an infection in the plant, which also requires certain virulence factors. On the contrary, during the CAT fusion, stress response genes were expressed so as to cope up with the stress. Further, in order to form CAT, an extensive transport, signalling and cell wall remodelling is required therefore genes coding for these processes were found be highly expressed during CAT fusion. Since CAT fusion involved formation of a bridging tube and transfer of cell organelles including nuclei, many genes coding for cytoskeleton and cell cycle regulation are expressed uniquely during CAT fusion. Our study demonstrates mutual exclusiveness of GT formation versus CAT fusion and also deciphered unique molecular requirements for these two processes in *C. gloeosporioides*. It would be interesting to know whether other fungal species capable of forming CAT also exhibit similar transcriptome profiles.

**Fig. 9:**
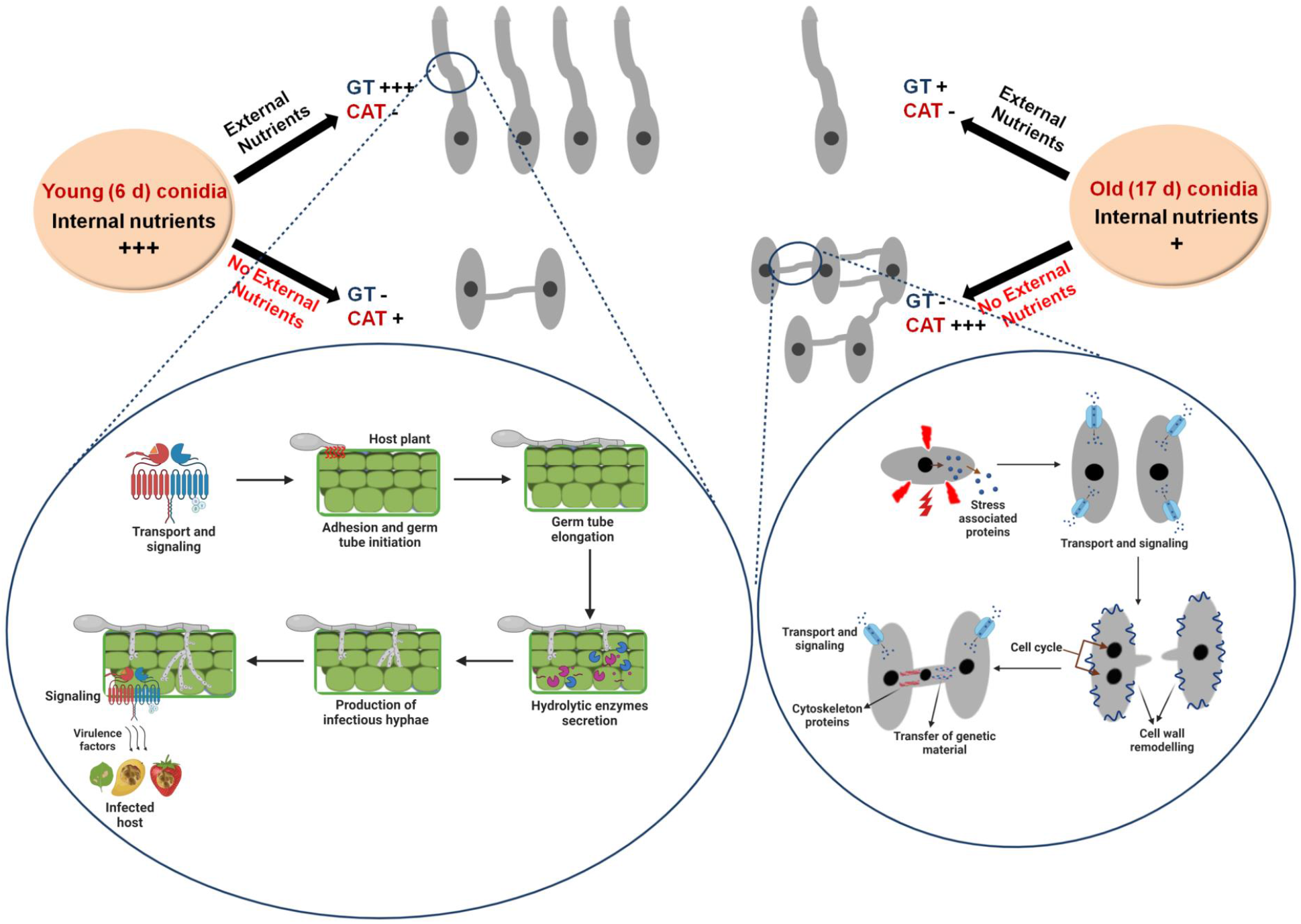
Model representing the differential physiological and molecular requirements of GT formation and CAT fusion in *C. gloeosporioides*.

## Author contributions

NM and AB designed the research, NM performed the experiments, NM, RP and AB analyzed and interpreted the experiments and results. They also wrote the manuscript; all authors read, revised and approved the manuscript.

## Acknowledgements

We are thankful to the Director, MACS’ Agharkar Research Institute for providing the necessary facility to carry out the research work. AB is thankful to SERB-DST for sanctioning the core research grant (CRG/2018/001786). NM acknowledges Council for Scientific and Industrial Research (CSIR) New Delhi for the senior research fellowship.

