## Supplementary file for "Deciphering the differential physiological and molecular requirements for conidial anastomosis tube fusion and germ tube formation in *Colletotrichum gloeosporioides*"

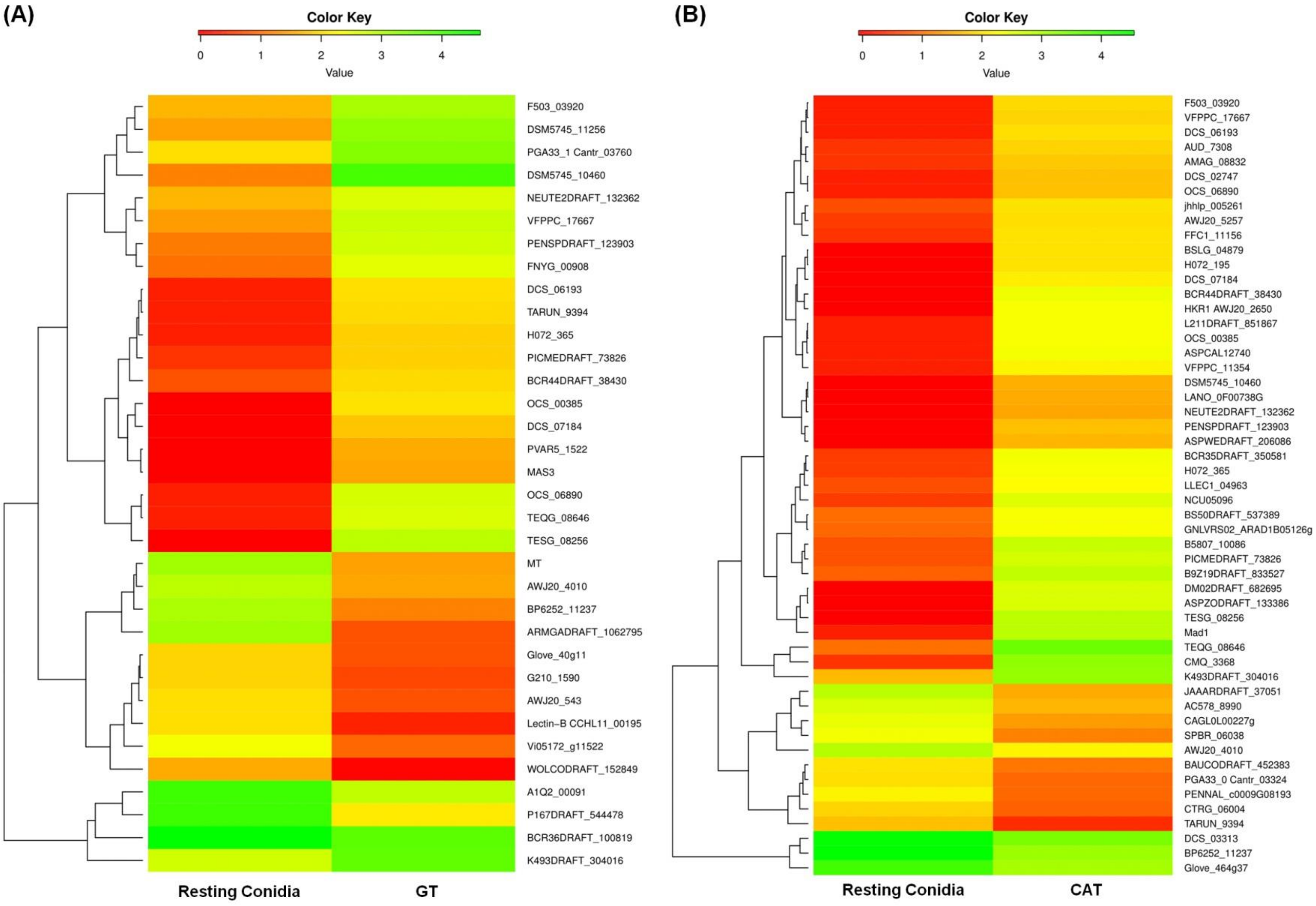


**Fig. S1: Heatmaps of differential expression profile of selected genes between A)** resting conidia versus GT formation. **B)** resting conidia versus CAT fusion.

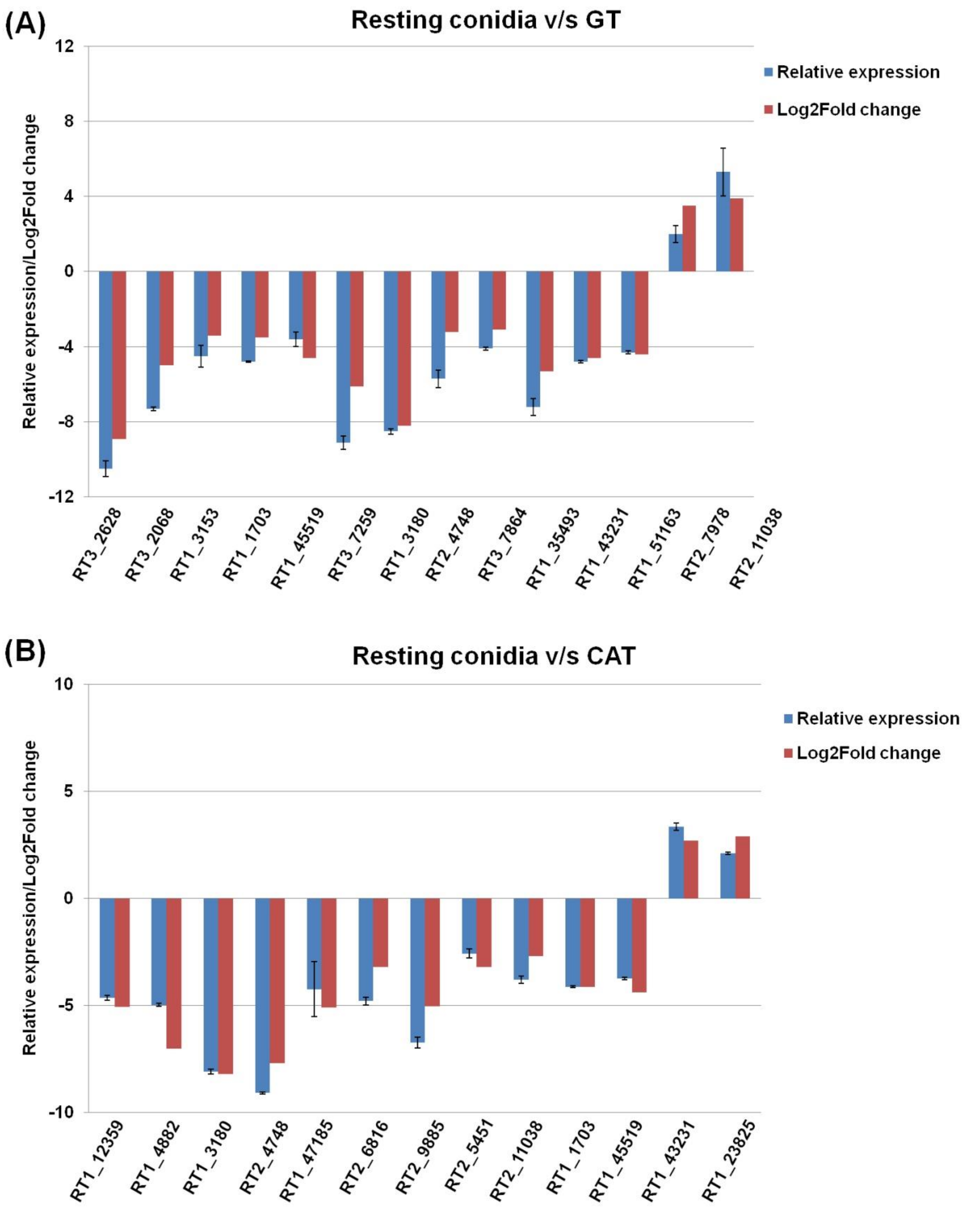


**Fig. S2: A graph of the level of correlation between relative expression obtained by qRT-PCR and RNA-Seq log2fold change of A)** 14 DEGs in resting conidia versus GT formation. **B)** 13 DEGs in resting conidia versus CAT fusion.

**Table S1**: Read statistics of prepared libraries of resting conidia, GT and CAT.

| **Samples** | **Resting conidia** | **GT** | **CAT** |
|---|---|---|---|
| Raw Reads | 49196663 | 35626747 | 40063840 |
| Processed Reads | 46579519 | 33940707 | 38264066 |
| Alignment to clustered transcripts (%) | 95.80% | 94.97% | 95.47% |

**Table S2:** Assembly statistics of prepared libraries of resting conidia, GT and CAT.

| | **Resting conidia** | **GT** | **CAT** |
|---|---|---|---|
| Number of clustered transcripts | 62064 | 55736 | 65536 |
| Maximum Contig Length | 30968 | 31011 | 24379 |
| Minimum Contig Length | 31 | 42 | 36 |
| Average Contig Length | 663.5 | 667.4 | 608.3 |
| Median Contig Length | 253 | 260 | 250 |
| Total Contigs Length | 41182192 | 37196863 | 39867188 |
| Total Number of Non-ATGC Characters | 106913 | 73863 | 89018 |
| Contigs >= 100 bp | 61715 | 55504 | 65223 |
| Contigs >= 200 bp | 46451 | 43084 | 47931 |
| Contigs >= 500 bp | 14056 | 13795 | 13542 |
| Contigs >= 1 Kbp | 9007 | 8826 | 8415 |
| Contigs >= 10 Kbp | 98 | 38 | 81 |
| Contigs >= 1 Mbp | 0 | 0 | 0 |
| N50 value | 2071 | 1880 | 1829 |

| **Combined assembly statistics** | |
|---|---|
| Number of clustered transcripts | 121260 |
| Maximum Contig Length | 31011 |
| Minimum Contig Length | 37 |
| Average Contig Length | 603.6 |
| Median Contig Length | 274 |
| Total Contigs Length | 73195249 |
| Total Number of Non-ATGC Characters | 233311 |
| Contigs >= 100 bp | 121063 |
| Contigs >= 200 bp | 97616 |
| Contigs >= 500 bp | 22246 |
| Contigs >= 1 Kbp | 14404 |
| Contigs >= 10 Kbp | 173 |
| Contigs >= 1 Mbp | 0 |
| N50 value | 1849 |

**Table S3:** Top 50 up-regulated genes during GT formation.

| Transcript ID | GT Expression | CAT Expression | log2 Fold Change | P value | Gene ID | Gene name | Possible functions |
|---|---|---|---|---|---|---|---|
| RT2_5451 | 1712 | 2 | 9.741466986 | 0.0001363 | CGLO_05791 | Pectin lyase | Degrades pectin (found in plant cell wall) |
| RT2_6913 | 1571 | 2 | 9.617467465 | 0.0001623 | CGGC5_6920 | Fg-gap repeat protein | Important for ligand binding |
| RT2_4144 | 1556 | 2 | 9.603626345 | 0.0001655 | CGGC5_6648 | Phosphate-repressible phosphate permease/ transporter | Membrane transport proteins that facilitate the diffusion of phosphate into and out of a cell or organelle |
| RT2_16712 | 761 | 1 | 9.571752644 | 0.0003735 | CGGC5_12869 | Quinone reductase | Host-fungus interaction & infection |
| RT2_3663 | 4430 | 6 | 9.528128483 | 0.0000810 | CGLO_01772 | Peptidase S41 family protein | Role in pathogenesis |
| RT1_10305 | 9563 | 16 | 9.22324756 | 0.0000912 | CGGC5_8072 | Hydrophobin 2 | Involved in adhesion |
| RT2_4587 | 3ed | 6 | 9.148900219 | 0.0001324 | CGLO_15861 | Transmembrane amino acid transporter | Promoting ubiquitin dependent endocytosis and signaling responses |
| RT2_7597 | 1122 | 2 | 9.131856961 | 0.0003204 | CGLO_03631 | Short-chain dehydrogenase/reductase ATR9 | Involved in Mycotoxin biosynthesis |
| RT3_10618 | 6328 | 13 | 8.927094166 | 0.0001330 | NA | Glycosyl hydrolase family 114 protein | Degrades cellulase, hemicellulose and lignin found in plant cell wall |
| RT2_4888 | 2852 | 6 | 8.892795766 | 0.0001857 | CGGC5_4502 | Extracellular solute-binding protein family 1 | Role in membrane transport |
| RT1_10110 | 6594 | 14 | 8.87958325 | 0.0001388 | NA | Glycosyl hydrolase family 114 protein | Degrades cellulase, hemicellulose and lignin found in plant cell wall |
| RT2_973 | 5602 | 12 | 8.866763767 | 0.0001459 | CGLO_08686 | Ferric reductase transmembrane component 4 | Iron acquisition and virulence |
| RT2_27038 | 1299 | 3 | 8.758223215 | 0.0003678 | CGGC5_3625 | Nonribosomal peptide synthetases (nrps)-like protein | Involved in virulence |
| RT2_7032 | 814 | 2 | 8.668884984 | 0.0005890 | CGLO_05279 | CHRD superfamily protein | Involved in the chemical reactions and pathways resulting in the breakdown of organonitrogen compound |
| RT2_3495 | 3561 | 10 | 8.476138625 | 0.0002533 | CGLO_11758 | Polysaccharide | Involved in virulence |

| | | | | | | | |
|---|---|---|---|---|---|---|---|
| | | | | | | deacetylase | |
| RT2_4111 | 7920 | 24 | 8.366322214 | 0.0002378 | CGLO_16353 | Glycosyl hydrolase family 76 | Degrades cellulase, hemicellulose and lignin found in plant cell wall |
| RT2_10908 | 625 | 2 | 8.28771238 | 0.0009541 | CGLO_09660 | FGGY-family pentulose kinase | Involved in virulence |
| RT2_10645 | 620 | 2 | 8.276124405 | 0.0009681 | CGGC5_10265 | Sodium chloride dependent neurotransmitter | Role in signaling |
| RT2_7569 | 589 | 2 | 8.202123824 | 0.0010620 | CGLO_03308 | Sodium:neurotransmitter symporter family protein | Role in signaling |
| RT2_3998 | 1738 | 6 | 8.178249866 | 0.0004869 | CGLO_05247 | Copper amine oxidase | Cell differentiation, growth, wound healing, detoxification and cell signalling |
| RT1_5572 | 17456 | 64 | 8.091435386 | 0.0003357 | CGGC5_12259 | Acetylornithine aminotransferase | Arginine biosynthesis |
| RT1_4917 | 16867 | 62 | 8.087719465 | 0.0003354 | CGLO_14124 | Cerato-platanin | Role in fungus-plant interactions and infection |
| RT2_4630 | 12530 | 47 | 8.058509943 | 0.0003358 | CGLO_03526 | Aminotransferase class-III | Regulate the fungus-host interaction |
| RT1_11723 | 3726 | 14 | 8.056057037 | 0.0003870 | CGLO_09660 | FGGY-family pentulose kinase | Involved in virulence |
| RT2_5343 | 488 | 2 | 7.930737338 | 0.0014906 | CGLO_15345 | Histidine acid phosphatase | Roles in cellular regulation and signaling |
| RT2_2831 | 1334 | 6 | 7.79658045 | 0.0008204 | CGGC5_4931 | Amino acid transporter | Promoting ubiquitin dependent endocytosis and signaling responses |
| RT2_7681 | 3090 | 14 | 7.786036201 | 0.0005480 | CGGC5_1286 | Spherulin 4-like cell surface | Involved in virulence |
| RT2_3794 | 13395 | 67 | 7.64331777 | 0.0005425 | CGGC5_14688 | Copper amine oxidase | Involved in cell differentiation and growth, wound healing, detoxification and cell signaling |
| RT2_12884 | 190 | 1 | 7.569855608 | 0.0049030 | CGGC5_24 | Het-s domain protein | HET Domain Mediates Programmed Cell Death |
| RT1_15737 | 182 | 1 | 7.50779464 | 0.0052847 | CGGC5_4042 | MFS monosaccharide transporter | Membrane transport proteins |
| RT2_7093 | 362 | 2 | 7.499845887 | 0.0025589 | CGLO_16009 | Methyltransferase domain-containing protein | Epigenetic regulation of fungal development |

| | | | | | | | |
|---|---|---|---|---|---|---|---|
| RT2_8698 | 347 | 2 | 7.438791853 | 0.0027641 | CGGC5_14315 | Bifunctional pyrimidine biosynthesis protein | Involved in pyrimidine biosynthesis |
| RT2_4766 | 1027 | 6 | 7.419257966 | 0.0013549 | CGGC5_8513 | High affinity methionine permease | Cysteine and methionine transport across plasma membrane |
| RT2_5240 | 324 | 2 | 7.339850003 | 0.0031333 | CGLO_12005 | GMC oxidoreductase | Role in pathogenicity |
| RT1_5191 | 12105 | 77 | 7.296528918 | 0.0008012 | CGLO_15244 | Beta-glucuronidase | Members of the glycosyl hydrolase family, role in degradation of plant cell wall |
| RT2_9014 | 934 | 6 | 7.282316239 | 0.0016145 | CGLO_02683 | Zinc RING finger of MSL2 | Exhibits ubiquitin E3 ligase activity, required for fungal virulence |
| RT2_2131 | 10433 | 68 | 7.261403602 | 0.0008282 | CGGC5_232 | GNAT family acetyltransferase | Role in chitin metabolism of fungi and is required for development and pathogenicity |
| RT2_5893 | 21332 | 143 | 7.220860276 | 0.0009399 | CGLO_01146 | GPI-anchored cell wall beta-1,3-endoglucanase EglC | Role in germination |
| RT2_7208 | 594 | 4 | 7.214319121 | 0.0021990 | CGGC5_10072 | Pectin lyase | Degrades pectin (found in plant cell wall) |
| RT2_6645 | 885 | 6 | 7.204571144 | 0.0017812 | NA | Peptidase C50 domain-containing protein | Role in pathogenesis |
| RT2_3394 | 587 | 4 | 7.197216693 | 0.0022457 | CGGC5_9379 | Peptidase s41 family protein | Role in pathogenesis |
| RT2_6404 | 1848 | 13 | 7.151309323 | 0.0012963 | CGLO_14363 | Mandelate racemase/muconate lactonizing enzyme domain-containing protein | Both are involved in the breakdown of lignin-derived aromatics |
| RT3_9669 | 24227 | 173 | 7.129699921 | 0.0010700 | CGLO_15642 | Uso1 / p115 like vesicle tethering protein | Intracellular protein transport |
| RT3_7789 | 86745 | 621 | 7.126043524 | 0.0016494 | CGLO_10547 | Hydrophobic surface binding protein A | Increase the hydrophobicity of conidia, aerial hyphae and fruiting bodies |
| RT2_7531 | 13700 | 100 | 7.098032083 | 0.0010163 | CGLO_08541 | CAP-22 protein | Expressed in the conidium during the appressorium formation |
| RT1_7969 | 4225 | 33 | 7.000341507 | 0.0011918 | CGGC5_7497 | NmrA family transcriptional regulator | nmrA is required for the invasive virulence |

| | | | | | | | |
|---|---|---|---|---|---|---|---|
| RT2_7311 | 7926 | 62 | 6.998180942 | 0.0011195 | CGLO_13134 | Complex I intermediate-associated protein 30 | Increased susceptibility to oxidative stress and phleomycin |
| RT2_6816 | 254 | 2 | 6.988684687 | 0.0049110 | CGLO_11115 | Fungal cellulose binding domain-containing protein | Plant cell-wall hydrolysis |
| RT1_7368 | 2902 | 23 | 6.979269848 | 0.0013290 | CGGC5_3105 | Cyanamide hydratase | Cyanamide hydratase involved in the detoxification and/or utilization of cyanamide |
| RT2_7165 | 65187 | 532 | 6.937014226 | 0.0017935 | CGLO_10547 | Hydrophobic surface binding protein A | Increase the hydrophobicity of conidiospores, aerial hyphae and fruiting bodies |

**Table S4:** Top 50 up-regulated genes during CAT fusion.

| Transcript ID | GT Expression | CAT Expression | log2 Fold Change | P value | Gene ID | Gene name | Possible functions |
|---|---|---|---|---|---|---|---|
| RT1_11804 | 2 | 14522 | 12.82595254 | 0.00000289 | CGLO_12901 | Major facilitator superfamily transporter | Membrane transport proteins |
| RT1_6087 | 2 | 8376 | 12.03204573 | 0.00000685 | CGGC5_7519 | Taurine catabolism dioxygenase | Role in utilisation of taurine under sulfate starvation |
| RT1_4872 | 2 | 5165 | 11.33455263 | 0.00001597 | CGGC5_4913 | Enoyl-hydratase isomerase family | Role in virulence |
| RT1_10064 | 10 | 18539 | 10.85634771 | 0.00001617 | CGLO_12901 | Major facilitator superfamily transporter | Membrane transport proteins |
| RT1_7001 | 2 | 2907 | 10.50531536 | 0.00004744 | MreB_Mbl (PF06723) | MreB/Mbl protein | Constituents of the eukaryotic cytoskeleton (tubulin, actin) |
| RT3_33317 | 5 | 7093 | 10.47025214 | 0.00002721 | CGGC5_4914 | Pyridine nucleotide-disulfide | Involved in cellular oxidative stress response |
| RT1_4751 | 390 | 509554 | 10.3515452 | 0.00027369 | CGGC5_4914 | Pyridine nucleotide-disulfide | Involved in cellular oxidative stress response |
| RT2_20649 | 4 | 5205 | 10.34568245 | 0.00003512 | CGLO_12901 | Major facilitator superfamily transporter | Membrane transport proteins |
| RT3_14378 | 20 | 24057 | 10.23224103 | 0.00003274 | CGGC5_3890 | Formate dehydrogenase | Is an oxidoreductase |
| RT1_47325 | 1 | 1058 | 10.04712391 | 0.00019978 | CGLO_13066 | Sugar transporter | Membrane transport proteins |
| RT3_5595 | 10 | 10508 | 10.03727239 | 0.00003804 | CGGC5_7519 | Taurine catabolism dioxygenase | Role in utilisation of taurine under sulfate starvation |
| RT3_3888 | 6 | 5287 | 9.783271111 | 0.00005867 | CGLO_03583 | Oxalate decarboxylase family bicupin | Associated with pathogenesis in plants |
| RT3_49072 | 2 | 1466 | 9.517669388 | 0.00018689 | LSU_rRNA_eukarya | Eukaryotic large subunit ribosomal RNA | Role in maintenance of diploidy in fungi, regulates centrosome duplication |
| RT3_26602 | 8 | 5805 | 9.503080352 | 0.00007475 | CGLO_17005 | NADH dehydrogenase | Involved in cellular oxidative stress response |
| RT3_59288 | 2 | 1332 | 9.379378367 | 0.00022730 | CGGC5_4147 | F-box domain protein | Involved in control of the cell division cycle, glucose sensing, mitochondrial connectivity, stress response |

| | | | | | | | |
|---|---|---|---|---|---|---|---|
| | | | | | | | and pathogenicity |
| RT1_7850 | 276 | 174041 | 9.300543229 | 0.00026448 | CGLO_02777 | Major facilitator superfamily transporter | Membrane transport proteins |
| RT3_2628 | 136 | 84350 | 9.276637607 | 0.00016255 | CGGC5_12199 | Dyp-type peroxidase family | Role in the degradation of lignin |
| RT3_43400 | 1 | 597 | 9.221587121 | 0.00058346 | CGGC5_3338 | DNA repair protein | Protects cells from DNA damage-induced genome instability and role in fungal virulence |
| RT1_6607 | 145 | 85105 | 9.197047184 | 0.00017795 | CGLO_02777 | Major facilitator superfamily transporter | Membrane transport proteins |
| RT3_1744 | 14 | 8132 | 9.182039578 | 0.00009697 | CGGC5_8447 | NADPH cytochrome p450 | Role in redox signaling |
| RT1_5411 | 17 | 8913 | 9.034232549 | 0.00011269 | CGLO_03583 | Oxalate decarboxylase family bicupin | Role in stress responses |
| RT1_47371 | 1 | 493 | 8.945443836 | 0.00082744 | CGLO_17434 | Major facilitator superfamily transporter | Membrane transport proteins |
| RT3_47963 | 3 | 1411 | 8.877539772 | 0.00031119 | CGLO_02777 | Major facilitator superfamily transporter | Membrane transport proteins |
| RT3_46229 | 5 | 2123 | 8.729960561 | 0.00025792 | CGLO_02777 | MFS multidrug transporter | Membrane transport proteins |
| RT3_34153 | 1 | 396 | 8.62935662 | 0.00123627 | NA | Lysosomal-associated transmembrane protein 4B | Promotes autophagy and cell survival under some stresses |
| RT1_52736 | 1 | 361 | 8.495855027 | 0.00146671 | CGGC5_7344 | Glutamyl-tRNA synthetase | Role in protein synthesis |
| RT1_45947 | 3 | 952 | 8.309855263 | 0.00067180 | CGLO_05737 | Cyanate hydratase | Used as Nitrogen source |
| RT2_50347 | 1 | 314 | 8.294620749 | 0.00190175 | CGLO_04105 | Dienelactone hydrolase | Dlh1 gene responsible for pathogenicity |
| RT1_35493 | 2 | 582 | 8.184875343 | 0.00108520 | CGLO_13144 | Zn 2cys6 transcription factor | Function in meiosis, regulation of genes involved in the stress response and pleiotropic drug resistance |
| RT3_2836 | 21 | 5699 | 8.084175655 | 0.00033899 | CGGC5_10167 | Carboxylic acid transport protein | Involved in pathogenesis |

| | | | | | | | |
|---|---|---|---|---|---|---|---|
| RT3_56167 | 2 | 538 | 8.071462363 | 0.00125042 | CGGC5_4914 | Pyridine nucleotide-disulfide | Involved in the cellular oxidative stress response |
| RT1_2744 | 21 | 5608 | 8.060953211 | 0.00034875 | CGGC5_10167 | Carboxylic acid transport protein | Involved in pathogenesis |
| RT3_3106 | 75 | 19971 | 8.056800263 | 0.00035741 | NA | Maleylacetate reductase | Catabolism of some usual aromatic compounds like quinol or resorcinol |
| RT2_1386 | 2703 | 717435 | 8.052142807 | 0.00281692 | CGGC5_14430 | MFS multidrug transporter | Membrane transport proteins |
| RT3_40341 | 2 | 524 | 8.033423002 | 0.00131125 | CGGC5_7427 | MFS monosaccharide transporter | Membrane transport proteins |
| RT3_7864 | 5 | 1297 | 8.019034669 | 0.00068399 | CGLO_04915 | DNA methyltransferases | Epigenetic regulation of fungal development |
| RT1_2872 | 71 | 17772 | 7.967571307 | 0.00038771 | NA | Tetraspanin 10 | Superfamily of small integral membrane proteins and required for pathogenicity |
| RT2_3974 | 86 | 21409 | 7.959665035 | 0.00040477 | CGLO_15466 | CorA-like Mg2+ transporter | Essential for growth, development and infection |
| RT1_50218 | 14 | 3253 | 7.860200185 | 0.00049785 | CGGC5_1275 | Sugar transporter | Membrane transport proteins |
| RT2_20124 | 2 | 429 | 7.744833837 | 0.00188076 | CGGC5_6603 | S-adenosylmethionine-dependent methyltransferase | Epigenetic regulation of fungal development |
| RT1_51991 | 1 | 213 | 7.73470962 | 0.00397874 | CGLO_03483 | Proline dehydrogenase | Role in osmotic, drought and salinity stress |
| RT3_8438 | 5 | 1046 | 7.708739041 | 0.00103544 | CGLO_01474 | Integral membrane protein | Required for pathogenicity |
| RT1_871 | 95 | 19654 | 7.692679732 | 0.00053989 | CGLO_11702 | Transmembrane amino acid transporter | Promots ubiquitin dependent endocytosis and signaling responses |
| RT1_45939 | 1 | 200 | 7.64385619 | 0.00447243 | CGGC5_7369 | ABC multidrug transporter | Multidrug resistance |
| RT1_3846 | 4 | 792 | 7.62935662 | 0.00131430 | CGLO_02115 | Integral membrane protein | Required for pathogenicity |
| RT1_2306 | 188 | 36701 | 7.608942901 | 0.00069427 | CGGC5_8255 | Phospholipid methyltransferase | Role in fungal development, fungicide resistance and virulence |

| | | | | | | | |
|---|---|---|---|---|---|---|---|
| RT3_50458 | 3 | 562 | 7.549463819 | 0.00175097 | CGLO_12789 | MutS domain V | DNA mismatch repair protein |
| RT3_56982 | 6 | 1118 | 7.541741972 | 0.00115507 | CGGC5_4147 | F-box domain protein | Involved in control of the cell division cycle, glucose sensing, mitochondrial connectivity, stress response and in pathogenicity |
| RT3_702 | 86 | 15743 | 7.516158113 | 0.00063860 | CGLO_11702 | Transmembrane amino acid transporter | Promots ubiquitin dependent endocytosis and signaling responses |
| RT1_52748 | 1 | 181 | 7.499845887 | 0.00533505 | CGGC5_9208 | Nitrate reductase | Required for nitrate uptake into fungal cells |

**Table S5:** Differentially expressed transcription factor proteins involved in GT formation and CAT fusion.

| S. N. | Protein names | Number of proteins | |
|---|---|---|---|
| | | GT | CAT |
| 1 | 2-dehydro-3-deoxygluconokinase (EC 2.7.1.45) | 1 | 0 |
| 2 | 2-isopropylmalate synthase | 0 | 1 |
| 3 | 30S ribosomal protein S12 | 1 | 4 |
| 4 | 3'-5'-exoribonuclease | 1 | 0 |
| 5 | 3-beta hydroxysteroid dehydrogenase/isomerase family protein | 21 | 16 |
| 6 | 3-demethylubiquinone-9 3-methyltransferase | 0 | 1 |
| 7 | 3-hydroxyacyl-CoA dehydrogenase, putative | 4 | 9 |
| 8 | 40S ribosomal protein S11-A (RP41) (S18) (Small ribosomal subunit protein uS17-A) (YS12) | 0 | 1 |
| 9 | 4-hydroxyphenylpyruvate dioxygenase | 0 | 1 |
| 10 | 50S ribosomal protein L19 | 0 | 1 |
| 11 | 54S ribosomal protein rml2, mitochondrial (L2) | 0 | 1 |
| 12 | 60S ribosomal protein L2 | 0 | 2 |
| 13 | 60S ribosomal protein L2-C (K37) (K5) (KD4) | 0 | 1 |
| 14 | Ab1-133 | 1 | 0 |
| 15 | ABC transporter ATP-binding protein (Duplicated ATPase domains) | 18 | 21 |
| 16 | Acetyl-CoA C-acetyltransferase | 0 | 1 |
| 17 | Acetyl-coenzyme A synthetase (Fragment) | 5 | 7 |
| 18 | Acyl-CoA dehydrogenase | 0 | 3 |
| 19 | Acyl-CoA dehydrogenase (Acyl-CoA dehydrogenase, N-terminal domain protein) (Butyryl-CoA dehydrogenase) (EC 1.3.8.1) | 12 | 20 |
| 20 | Acyl-CoA hydrolase (Acyl-CoA thioesterase) (Thioesterase superfamily protein) | 2 | 0 |
| 21 | Alcohol dehydrogenase (Cytochrome c) | 0 | 3 |
| 22 | Aliphatic sulfonates import ATP-binding protein SsuB (EC 3.6.3.-) | 0 | 2 |
| 23 | Alpha/beta hydrolase (Fragment) | 0 | 1 |
| 24 | Alpha-L-fucosidase | 1 | 3 |
| 25 | Arabinan endo-1,5-alpha-L-arabinosidase | 8 | 4 |
| 26 | Arginine--tRNA ligase (EC 6.1.1.19) (Arginyl-tRNA synthetase) (ArgRS) | 0 | 1 |
| 27 | Argininosuccinate synthase (EC 6.3.4.5) (Citrulline--aspartate ligase) | 3 | 6 |
| 28 | Asparagine-rich zinc-finger protein | 2 | 0 |
| 29 | Asparagine--tRNA ligase, cytoplasmic (EC 6.1.1.22) (Asparaginyl-tRNA synthetase) (AsnRS) | 0 | 1 |
| 30 | Aspartate--tRNA ligase, cytoplasmic (EC 6.1.1.12) (Aspartyl-tRNA synthetase) (AspRS) | 1 | 0 |
| 31 | Aspartate-tRNA(Asn) ligase | 1 | 0 |
| 32 | Aspartyl-tRNA synthetase | 8 | 13 |
| 33 | Aspyridones cluster regulator apdR (Aspyridones biosynthesis protein R) | 0 | 1 |
| 34 | ATPase (Fragment) | 0 | 1 |
| 35 | Binuclear zinc transcription factor | 11 | 6 |
| 36 | Branchpoint-bridging protein (Mud synthetic-lethal 5 protein) (Splicing | 0 | 1 |

| | | | |
|---|---|---|---|
| | factor 1) (Zinc finger protein BBP) | | |
| 37 | BZIP transcription factor | 2 | 5 |
| 38 | BZIP transcription factor (AtfA), putative | 0 | 4 |
| 39 | C2H2 zinc finger protein | 2 | 0 |
| 40 | C6 finger domain protein | 3 | 1 |
| 41 | C6 transcription factor (Gal4), putative | 0 | 2 |
| 42 | C6 transcription factor (UaY), putative | 1 | 0 |
| 43 | C6 transcription factor QutA, putative | 3 | 1 |
| 44 | C6 transcription factor RosA-like, putative | 0 | 1 |
| 45 | C6 transcription factor RosA | 0 | 1 |
| 46 | C6 transcription factor, putative | 123 | 138 |
| 47 | C6 zinc finger protein | 0 | 2 |
| 48 | Calcium-translocating P-type ATPase, PMCA-type | 19 | 24 |
| 49 | Carbohydrate kinase family protein (Fragment) | 0 | 1 |
| 50 | Carboxylic ester hydrolase (EC 3.1.1.-) | 56 | 44 |
| 51 | Casein kinase II subunit beta (CK II beta) | 0 | 1 |
| 52 | Cell pattern formation-associated protein stuA | 3 | 1 |
| 53 | Choline dehydrogenase (Fragment) | 22 | 12 |
| 54 | Chorismate synthase (CS) (EC 4.2.3.5) (5-enolpyruvylshikimate-3-phosphate phospholyase) | 1 | 4 |
| 55 | Class 2 transcription repressor NC2 | 0 | 2 |
| 56 | Conserved hypothetical membrane protein | 1 | 3 |
| 57 | Conserved hypothetical secreted protein | 0 | 1 |
| 58 | Cullin binding protein CanA, putative | 1 | 3 |
| 59 | Cullin-1 | 3 | 8 |
| 60 | Cutinase transcription factor 1 beta | 9 | 6 |
| 61 | Cyanate hydratase (Cyanase) (EC 4.2.1.104) (Cyanate hydrolase) (Cyanate lyase) | 1 | 3 |
| 62 | Cystathionine gamma-synthase (Fragment) | 9 | 4 |
| 63 | Cytochrome c oxidase polypeptide 4 (EC 1.9.3.1) (Cytochrome aa3 subunit 4) (Cytochrome c oxidase polypeptide IV) | 0 | 1 |
| 64 | Cytoplasmic 60S subunit biogenesis factor REI1 (Required for isotropic bud growth protein 1) (pre-60S factor REI1) | 1 | 0 |
| 65 | Cytoplasmic asparaginyl-tRNA synthetase | 1 | 4 |
| 66 | DEHA2A12694p | 0 | 1 |
| 67 | DEHA2D14718p | 0 | 1 |
| 68 | DEHA2E11682p | 0 | 4 |
| 69 | DEHA2G20328p | 0 | 1 |
| 70 | Deoxyribonuclease HsdR | 0 | 1 |
| 71 | Developmental regulator FlbA | 2 | 0 |
| 72 | Dihydroxyacetone kinase DhaK subunit | 5 | 3 |
| 73 | Dihydroxy-acid dehydratase (DAD) (EC 4.2.1.9) | 8 | 5 |
| 74 | DNA polymerase III subunit alpha (EC 2.7.7.7) | 1 | 5 |
| 75 | DNA replication licensing factor mcm2 (EC 3.6.4.12) (Cell division control protein 19) (Minichromosome maintenance protein 2) | 3 | 1 |
| 76 | DNA replication licensing factor mcm5 | 1 | 0 |
| 77 | DNA replication licensing factor MCM6 (EC 3.6.4.12) | 1 | 0 |

| | | | |
|---|---|---|---|
| | (Minichromosome maintenance protein 6) | | |
| 78 | DNA-binding protein HEXBP | 9 | 4 |
| 79 | Endo-1,4-beta-glucanase, putative | 8 | 11 |
| 80 | Eukaryotic translation initiation factor 5A-1 (eIF-5A-1) | 2 | 0 |
| 81 | Eukaryotic translation initiation factor 5A-2 (eIF-5A-2) | 0 | 2 |
| 82 | Exosome complex component rrp40 (Ribosomal RNA-processing protein 40) | 0 | 1 |
| 83 | Exosome complex exonuclease RRP4 | 3 | 1 |
| 84 | FAD-dependent oxidoreductase | 1 | 0 |
| 85 | Ferrous iron transport protein B | 0 | 1 |
| 86 | FKH1 transcription factor-like protein | 0 | 2 |
| 87 | Fructose-1,6-bisphosphate aldolase (Fructose-1,6-bisphosphate aldolase, class II) (Fructose-bisphosphate aldolase) (Fructose-bisphosphate aldolase class II) (EC 4.1.2.13) | 4 | 1 |
| 88 | Fungal specific transcription factor domain protein | 0 | 3 |
| 89 | Fused acetyl/propionyl-CoA carboxylase subunit alpha/methylmalonyl-CoA decarboxylase subunit alpha | 10 | 14 |
| 90 | Galactoside O-acetyltransferase | 4 | 2 |
| 91 | Gamma-glutamyltranspeptidase (EC 2.3.2.2) | 3 | 7 |
| 92 | Heat shock transcription factor | 0 | 1 |
| 93 | Heavy metal translocating P-type ATPase | 8 | 13 |
| 94 | Helicase | 8 | 11 |
| 95 | Helix-loop-helix DNA-binding domain-containing protein | 2 | 0 |
| 96 | High-affinity nicotinic acid transporter | 72 | 92 |
| 97 | High-copy mep suppressor | 0 | 1 |
| 98 | Homeobox domain-containing protein | 2 | 0 |
| 99 | Homeobox transcription factor phx1 | 0 | 1 |
| 100 | Homeobox transcription factor, putative | 3 | 0 |
| 101 | Homoserine kinase (HK) (HSK) (EC 2.7.1.39) | 1 | 0 |
| 102 | Hydrophobe/amphiphile efflux-1 (HAE1) family transporter | 0 | 2 |
| 103 | Imidazolonepropionase (EC 3.5.2.7) (Imidazolone-5-propionate hydrolase) | 0 | 3 |
| 104 | Inosine-5'-monophosphate dehydrogenase (IMP dehydrogenase) (IMPD) (IMPDH) (EC 1.1.1.205) | 2 | 6 |
| 105 | Involucrin repeat protein | 1 | 2 |
| 106 | Iron-sulfur protein | 1 | 0 |
| 107 | Isocitrate dehydrogenase [NADP] (EC 1.1.1.42) | 4 | 8 |
| 108 | J protein JJJ1 | 0 | 1 |
| 109 | Jmjc domain-containing histone demethylase | 0 | 1 |
| 110 | J-protein (Type III) | 0 | 1 |
| 111 | Kinesin-like protein bimC | 0 | 2 |
| 112 | Leptomycin B resistance protein pmd1 | 49 | 56 |
| 113 | Leucine dehydrogenase (EC 1.4.1.9) | 1 | 0 |
| 114 | Leucyl/phenylalanyl-tRNA--protein transferase (EC 2.3.2.6) (L/F-transferase) (Leucyltransferase) (Phenyalanyltransferase) | 0 | 1 |
| 115 | Lid2 complex component lid2 | 1 | 3 |
| 116 | Lysine biosynthesis regulatory protein LYS14 | 1 | 0 |

| | | | |
|---|---|---|---|
| 117 | Lysine--tRNA ligase (EC 6.1.1.6) (Lysyl-tRNA synthetase) | 11 | 15 |
| 118 | Malate dehydrogenase (EC 1.1.1.37) | 2 | 0 |
| 119 | MeaB protein | 1 | 3 |
| 120 | Membrane dipeptidase (Peptidase family M19) | 0 | 2 |
| 121 | Methionine gamma-lyase | 3 | 7 |
| 122 | Methionyl-tRNA formyltransferase (EC 2.1.2.9) | 5 | 1 |
| 123 | Methylmalonyl-CoA carboxyltransferase | 4 | 1 |
| 124 | Methylmalonyl-CoA mutase (Fragment) | 0 | 2 |
| 125 | Minichromosome maintenance-related protein | 1 | 0 |
| 126 | Mis12-Mtw1 family protein | 1 | 0 |
| 127 | MIZ zinc finger protein | 7 | 3 |
| 128 | Molecular chaperone DnaJ | 13 | 19 |
| 129 | Multiprotein-bridging factor 1 | 1 | 0 |
| 130 | MYB family conidiophore development protein FlbD | 0 | 1 |
| 131 | Myb-like DNA-binding domain protein | 0 | 1 |
| 132 | N-acetyldiaminopimelate deacetylase (EC 3.5.1.47) | 1 | 0 |
| 133 | Negative regulator of pleiotropic drug resistance STB5 | 0 | 1 |
| 134 | NF-X1 finger and helicase domain protein, putative | 2 | 7 |
| 135 | NF-X1 finger transcription factor, putative | 0 | 3 |
| 136 | Nicotinate dehydrogenase small FeS subunit (EC 1.17.1.5) | 0 | 3 |
| 137 | Nodulation protein L | 4 | 1 |
| 138 | Non-histone chromosomal protein | 1 | 0 |
| 139 | Nonribosomal peptide synthase, putative | 78 | 14 |
| 140 | Nucleic binding protein | 1 | 0 |
| 141 | Nucleolar protein NOP2 | 5 | 7 |
| 142 | O-acetylhomoserineaminocarboxypropyltransferase (EC 2.5.1.49) | 5 | 9 |
| 143 | Oligopeptide transporter family protein | 1 | 0 |
| 144 | O-methyltransferase | 9 | 6 |
| 145 | Origin recognition complex subunit 4 | 1 | 4 |
| 146 | Outer membrane efflux protein | 1 | 3 |
| 147 | Oxidoreductase | 23 | 25 |
| 148 | Oxidoreductase, 2OG-Fe(II) oxygenase family | 15 | 19 |
| 149 | Palmitoyltransferase (EC 2.3.1.225) | 22 | 29 |
| 150 | Pfs, NACHT and WD domain protein (EC 2.4.2.-) | 430 | 380 |
| 151 | Phosphate transport system permease protein | 0 | 1 |
| 152 | Phosphatidylserine decarboxylase proenzyme 2, mitochondrial (EC 4.1.1.65) [Cleaved into: Phosphatidylserine decarboxylase 2 beta chain; Phosphatidylserine decarboxylase 2 alpha chain] | 3 | 5 |
| 153 | Phosphonates import ATP-binding protein PhnC (EC 7.3.2.2) | 0 | 1 |
| 154 | Phosphoribosylaminoimidazolesuccinocarboxamide synthase (Fragment) | 1 | 0 |
| 155 | Polyadenylation factor subunit CstF64, putative | 0 | 1 |
| 156 | Polyribonucleotide nucleotidyltransferase (EC 2.7.7.8) (Polynucleotide phosphorylase) (PNPase) | 0 | 2 |
| 157 | Potential protein lysine methyltransferase SET5 (EC 2.1.1.-) (SET domain-containing protein 5) | 0 | 1 |
| 158 | Pre-mRNA-splicing ATP-dependent RNA helicase PRP28 | 15 | 26 |

| | | | |
|---|---|---|---|
| 159 | Probable E3 ubiquitin-protein ligase HUL4 (EC 2.3.2.26) (HECT ubiquitin ligase 4) (HECT-type E3 ubiquitin transferase HUL4) | 1 | 4 |
| 160 | Proteasome regulatory particle subunit Rpt6, putative | 16 | 19 |
| 161 | Proteasome-activating nucleotidase | 5 | 7 |
| 162 | PUT3-like fungal specific transcription factor, putative | 0 | 4 |
| 163 | Putative histone demethylase JARID1D | 0 | 2 |
| 164 | Putative methyl-accepting chemotaxis protein | 0 | 1 |
| 165 | Putative multiprotein-bridging factor 1 | 2 | 4 |
| 166 | Putative NAD(P) transhydrogenase alpha subunit (EC 1.6.1.2) | 1 | 0 |
| 167 | Pyrophosphate--fructose 6-phosphate 1-phosphotransferase (EC 2.7.1.90) (6-phosphofructokinase, pyrophosphate dependent) (PPi-dependent phosphofructokinase) (PPi-PFK) (Pyrophosphate-dependent 6-phosphofructose-1-kinase) | 1 | 0 |
| 168 | Quinic acid utilization activator | 1 | 0 |
| 169 | Regulatory protein alcR (Fragment) | 0 | 1 |
| 170 | Replication protein A subunit | 2 | 4 |
| 171 | Replicative DNA helicase (EC 3.6.4.12) | 0 | 1 |
| 172 | Retaining alpha-galactosidase (EC 3.2.1.22) | 0 | 1 |
| 173 | RhoGAP and Fes/CIP4 domain protein | 2 | 0 |
| 174 | Rho-GTPase-activating protein 8 | 1 | 3 |
| 175 | Riboflavin biosynthesis protein RibBA [Includes: 3,4-dihydroxy-2-butanone 4-phosphate synthase (DHBP synthase) (EC 4.1.99.12); GTP cyclohydrolase-2 (EC 3.5.4.25) (GTP cyclohydrolase II)] | 5 | 3 |
| 176 | Ribosomal protein S12 methylthiotransferase RimO (S12 MTTase) (S12 methylthiotransferase) (EC 2.8.4.4) (Ribosomal protein S12 (aspartate-C(3))-methylthiotransferase) (Ribosome maturation factor RimO) | 0 | 1 |
| 177 | Ribosomal protein S23 (S12) | 1 | 0 |
| 178 | Rubredoxin | 0 | 1 |
| 179 | Sensory transduction protein regX3 | 1 | 2 |
| 180 | Short chain dehydrogenase | 6 | 9 |
| 181 | Short-chain dehydrogenase, putative | 3 | 6 |
| 182 | Siderophore transcription factor SreA | 4 | 2 |
| 183 | Sigma-54 modulation protein | 0 | 1 |
| 184 | Soluble hydrogenase 42 kDa subunit (EC 1.12.-.-) | 1 | 3 |
| 185 | Specific RNA polymerase II transcription factor | 0 | 2 |
| 186 | Stage IV sporulation protein A (EC 3.6.1.3) (Coat morphogenetic protein SpoIVA) | 0 | 1 |
| 187 | Tagatose-6-phosphate kinase (EC 2.7.1.144) | 0 | 1 |
| 188 | telomere-associated protein 1 | 0 | 1 |
| 189 | TetR family transcriptional regulator | 0 | 2 |
| 190 | Thiamine repressible genes regulatory protein thi5 (Transcription factor ntf1 5) | 1 | 0 |
| 191 | Thiamine ABC transporter permease (Fragment) | 0 | 5 |
| 192 | TonB-dependent outer membrane receptor | 0 | 1 |
| 193 | Transcription activator of gluconeogenesis acuK (Acetate non-utilizing mutant protein K) | 0 | 1 |

| | | | |
|---|---|---|---|
| 194 | Transcription activator of gluconeogenesis BDCG_02812 | 0 | 1 |
| 195 | Transcription elongation factor GreA (Transcript cleavage factor GreA) | 0 | 1 |
| 196 | Transcription factor AbaA | 1 | 0 |
| 197 | Transcription regulatory protein SNF2 (EC 3.6.4.-) (ATP-dependent helicase SNF2) (Regulatory protein GAM1) (Regulatory protein SWI2) (SWI/SNF complex component SNF2) (Transcription factor TYE3) | 2 | 0 |
| 198 | Transcriptional activator hac1 | 2 | 5 |
| 199 | Transcriptional activator HAP2 | 0 | 1 |
| 200 | Transcriptional activator of proteases prtT (Zn(2)-C6 zinc finger-containing protein prtT) | 3 | 0 |
| 201 | Transcriptional activator protein acu-15 | 1 | 5 |
| 202 | Transcriptional activator xlnR | 1 | 4 |
| 203 | Transport system permease protein | 0 | 2 |
| 204 | tRNA dimethylallyltransferase (EC 2.5.1.75) | 3 | 1 |
| 205 | tRNA pseudouridine synthase B (EC 5.4.99.25) (tRNA pseudouridine(55) synthase) (Psi55 synthase) (tRNA pseudouridylate synthase) (tRNA-uridine isomerase) | 2 | 4 |
| 206 | Two-component system response regulator | 2 | 6 |
| 207 | Ubiquitin ligase subunit CulD | 2 | 6 |
| 208 | Ubiquitinyl hydrolase 1 (EC 3.4.19.12) | 6 | 4 |
| 209 | Uncharacterized FAD-linked oxidoreductase yvdP (EC 1.21.-.-) | 28 | 20 |
| 210 | UPF0276 protein BN444_02599 | 0 | 1 |
| 211 | UV radiation resistance protein (UVRAG), putative | 4 | 2 |
| 212 | Vacuolar segregation protein PEP7 | 1 | 0 |
| 213 | Voltage-gated chloride channel (ClcA), putative | 8 | 14 |
| 214 | Zinc finger protein 32 | 8 | 3 |
| 215 | Zinc finger protein 58 | 13 | 24 |
| 216 | Zinc finger protein GIS2 | 6 | 9 |
| 217 | Zinc knuckle domain containing protein | 3 | 6 |
| 218 | Zinc knuckle transcription factor (CnjB) | 2 | 6 |
| 219 | Zinc-responsiveness transcriptional activator | 4 | 0 |
| 220 | Zn(II)2Cys6 transcription factor | 2 | 4 |

**Table S6:** Uniquely expressed effector candidate genes during GT formation.

| Transcript_ID | Gene names | Possible functions |
|---|---|---|
| RT2_3129.p1 | 3-phytase | Hydrolytic enzyme acting on phosphoric monoester bonds |
| RT2_4851.p1 | 5'-nucleotidase, C-terminal domain | Hydrolytic enzyme, catalyzes the phosphorylytic cleavage of 5'nucleotides |
| RT2_21460.p1 | ABC transporter | Involved in appresorium formation |
| RT2_4955.p1 | Acetyl esterase | Hydrolytic enzyme, role in degradation of hemicelluloses and pectin |
| RT2_7714.p1 | Acetylcholinesterase | Catalyzes the breakdown of acetylcholine |
| RT2_11269.p2 | Acid phosphatase, putative | Role in the metabolic process of fungus growth |
| RT2_1963.p1 | Acid trehalase | It can furnish endogenous carbon and energy to the cell during germination of spores |
| RT2_12928.p2 | Alkaline phosphatase | Role in phosphate mobilisation from organic substrates under phosphate starvation |
| RT2_17326.p1 | Allantoinase | Has hydrolase activity |
| RT2_816.p1 | Alpha-mannosidase family protein | Role in host colonization |
| RT2_9649.p1 | Amidohydrolase | Type of hydrolase that acts upon amide bonds. |
| RT2_12914.p1 | Amino acid permease | Role in amino acid transport across plasma membrane. |
| RT2_6465.p1 | Aminotransferase class I and II | Regulate the fungus-host interaction |
| RT2_7682.p1 | Ankyrin repeat domain protein | Plays a role in both disease resistance and antioxidation metabolism |
| RT2_13205.p1 | Arginine deiminase type-3 | Might play role in acid resistance and intracellular survival |
| RT2_6250.p1 | Arsenate reductase | Role in arsenic Resistance |
| RT2_21406.p1 | ATP synthase F0 | Role in virulence |
| RT2_12723.p2 | Bacterial extracellular solute-binding protein, family 3 | Serve as chemoreceptors and initiators of signal transduction pathways and transport |
| RT2_12221.p1 | Bacteriodes thetaiotaomicron symbiotic chitinase | Induced the degradation of fungal cell wall, mediated via chitinase-like activity |
| RT2_4824.p1 | Berberine bridge enzyme | Is a cellobiose oxidase |

| | | |
|---|---|---|
| RT2_14227.p1 | Beta-1,3-endoglucanase | Play a major role in cell wall softening |
| RT2_4851.p1 | Calcineurin-like phosphoesterase | It governs stress survival, sexual differentiation and virulence |
| RT2_6726.p1 | Carbonic anhydrase | Role in fruiting body development and ascospore germination |
| RT2_6589.p1 | Catalase-peroxidase | Required for virulence |
| RT2_3.p1 | Cell agglutination protein mam3 | Involved in agglutination during conjugation |
| RT2_3176.p1 | Chitin recognition protein | Role in fungus-host interaction |
| RT2_7032.p1 | CHRD domain | Involved in the chemical reactions and pathways resulting in the breakdown of organonitrogen compound |
| RT2_4368.p1 | Class III aminotransferase, putative | Regulate the fungus-host interaction |
| RT2_14681.p1 | Coagulation factor 5/8 type domain-containing protein | Involved in cell adhesion |
| RT2_1622.p1 | Copper fist dna binding domain-containing protein (Fragment) | Role in pathogenicity |
| RT2_5487.p1 | Covalently-linked cell wall protein | Role in fitness and virulence |
| RT2_5938.p1 | Developmentally Regulated MAPK Interacting protein | Role in cell wall integrity pathway, diverse stresses and developmental processes |
| RT2_6828.p1 | Dienelactone hydrolase | Responsible for pathogenicity |
| RT2_10729.p1 | Endo-chitosanase | Role in degradation of the fungal cell wall |
| RT2_2892.p2 | Endoglucanase 3 | Role in germination |
| RT2_19796.p1 | Endomembrane protein 70 | Role in cellular adhesion, filamentous growth and endosome-to-vacuole sorting |
| RT2_7832.p1 | Endopeptidase | Role in pathogenesis |
| RT2_18223.p1 | Eukaryotic porin | It is an important regulator of $Ca^{2+}$ transport in and out of the mitochondria |
| RT2_3766.p1 | Exosome complex | Involved in transcript/mRNA processing |

| | | |
|---|---|---|
| | exonuclease RRP43 | |
| RT2_9596.p1 | Extracellular cellulase | Degrades cellulose and some other related polysaccharides |
| RT2_1608.p1 | FAD binding domain | Play a vital role in energy transfer and utilization during fungal growth and mycelia aggregation |
| RT2_8805.p1 | FAD dependent oxidoreductase | Required for biosynthesis of secondary metabolites resulting in virulence |
| RT2_9375.p1 | FAD-dependent oxygenase | Involved in cell wall biogenesis and melanization processes |
| RT2_3702.p1 | F-box domain protein | Involved in control of the cell division cycle, glucose sensing, mitochondrial connectivity, stress response and in pathogenicity |
| RT2_10852.p2 | Fungal hydrophobin | Hydrophobins mediate hyphal attachment to hydrophobic surfaces such as those of plants |
| RT2_14634.p1 | Fungal specific transcription factor domain-containing protein | Function in various cellular process like growth, survival, stress response |
| RT2_618.p1 | Galactose mutarotase-like | Involved in D-galactose utilization |
| RT2_3667.p1 | Galactose oxidase | They catalyzes the oxidation of a range of primary alcohols to the corresponding aldehyde |
| RT2_4717.p1 | GEgh16 protein | Required for appressorium formation |
| RT2_241.p1 | Glycoprotein x | Involved in infection |
| RT2_1942.p1 | Glycosyl hydrolase family 15 | Degrades cellulase, hemicellulose and lignin found in plant cell wall |
| RT2_15512.p2 | Glycosyl hydrolase family 28 | Degrades cellulase, hemicellulose and lignin found in plant cell wall |
| RT2_618.p1 | Glycosyl hydrolase family 31 | Degrades cellulase, hemicellulose and lignin found in plant cell wall |
| RT2_5434.p1 | Glycosyltransferase sugar-binding region containing DXD motif | Pathogenesis of plants by enabling hyphal growth |
| RT2_19314.p1 | HET domain-containing protein | HET domain mediates programmed cell death |
| RT2_16981.p1 | Hexose transporter | Transport glucose as well as plant cell wall derived sugars |
| RT2_10358.p1 | Hydrophobin | Hydrophobins mediate hyphal attachment to hydrophobic surfaces such as those of plants |

| | | |
|---|---|---|
| RT2_10252.p1 | Idi-2 | Function unknown or may be involved in programmed cell death |
| RT2_17444.p1 | Ketoacyl-synthetase C-terminal extension | Involved in fatty acid synthesis pathway |
| RT2_8244.p1 | L-amino-acid oxidase 2 | Play a wide range of biological functions either in basal amino acid catabolism or in reactions related to generation of $H_2O_2$ |
| RT2_10013.p1 | Leucine rich repeat N-terminal domain | Required for cAMP signaling, hyphal growth and virulence |
| RT2_10136.p1 | Lipocalin-like domain | Role in transportation of pheromones and small hydrophobic molecules such as steroids, bilins, retinoids and lipids |
| RT2_19572.p3 | Lysine-specific metallo-endopeptidase (MEP) | Binds strongly to beta-1,3-glucan and chitin, major polysaccharides constituting the fungal cell wall |
| RT2_9596.p1 | Lytic transglycolase | Role in cell morphology and virulence |
| RT2_4975.p1 | Manganese lipoxygenase | Accelerates programmed spore germination |
| RT2_3955.p1 | Melibiase | Responsible for virulence |
| RT2_6749.p1 | MMC protein (Fragment) | Involved in microcycle conidiation which is a survival mechanism for some fungi encountering unfavorable conditions, in which asexual spores germinate secondary spores directly without formation of mycelium. |
| RT2_16681.p1 | Monooxygenase | Play diverse and pivotal roles in versatile metabolism and fungal adaptation to specific ecological niches |
| RT2_10314.p2 | Nad-dependent 15-hydroxyprostaglandin dehydrogenase | An oxidoreductase |
| RT2_20272.p1 | NADPH--cytochrome p450 reductase | Involved in redox signaling |
| RT2_8448.p2 | Necrosis inducing protein | Role in infection |
| RT2_618.p1 | N-terminal barrel of NtMGAM and CtMGAM, maltase-glucoamylase | Responsible for hydrolysis of starch into polymers resulting in degradation of plant cell wall |

| | | |
|---|---|---|
| RT2_20121.p1 | OST3 / OST6 family, transporter family | Tolerance to abiotic stresses |
| RT2_13845.p2 | Oxidoreductase | Role in pathogenicity |
| RT2_4942.p1 | Peptidase m14 | Role in pathogenesis |
| RT2_1608.p1 | Phenol hydroxylase, C-terminal dimerisation domain | Induce melanization of fungi and contribute to enhanced virulence and fungal resistance to diverse environments |
| RT2_11269.p2 | Phosphoinositide phospholipase C, Ca2+-dependent | Play multiple cellular roles including growth, stress tolerance, sexual development, and virulence in fungi |
| RT2_13809.p1 | Pollen proteins Ole e I like | Might play a role in germination |
| RT2_825.p1 | Pro-kumamolisin | Members of this family are found in various subtilase propeptides |
| RT2_4018.p1 | Proteasome A-type and B-type | Required for fungal pathogenicity |
| RT2_3209.p1 | Putative glycosyl hydrolase family 3 N terminal domain-containing protein | Degrades cellulase, hemicellulose and lignin found in plant cell wall |
| RT2_11973.p1 | Putative GTPase activating protein for Arf | Regulate membrane/protein trafficking, filamentous growth, cell wall integrity and virulence |
| RT2_4014.p1 | Putative stress-responsive nuclear envelope protein | Role in maintaining cell viability during stationary phase induced by stress response |
| RT2_10571.p1 | Related to glycine-rich RNA-binding protein | Regulates virulence, development and stress responses |
| RT2_4412.p1 | Restculine oxidase | Has an oxidoreductase activity |
| RT2_7683.p2 | Rhomboid family | Involved in hypoxia sensing, signalling and virulence |
| RT2_11292.p1 | Sarcosine oxidase | Role in polarized growth and conidiation |
| RT2_3804.p1 | Secretory lipase, putative | Potential virulence factors |
| RT2_15444.p1 | Serum amyloid A protein | Role in host-fungus interaction during fungal infection |
| RT2_9393.p1 | Sphingomyelin | Associated with some pathogenic mechanisms |

| | | |
|---|---|---|
| | phosphodiesterase | |
| RT2_18566.p1 | Spindle poison sensitivity protein | This protein is associated with microtubule formation |
| RT2_4014.p1 | Stress response protein | Important in fungal adaptation to a wide range of stress conditions |
| RT2_19796.p1 | Transmembrane 9 superfamily member | Essential for cell adhesion and filamentous growth |
| RT2_9804.p1 | Trypsin and protease inhibitor | Important in defense mechanism |
| RT2_6972.p1 | Tyrosinase central domain protein | Involved in melanin production |
| RT2_7974.p1 | Ubiquitin 3 binding protein But2 C-terminal domain | Required for fungal pathogenicity |
| RT2_156.p1 | Udp-glucose:glycoprotein glucosyltransferase | Role in growth and development |
| RT2_5435.p1 | Versicolorin b synthase | Involved in biosynthesis of aflatoxins |
| RT2_10806.p1 | WD40 domain protein beta propeller | Regulates fungal cell differentiation |
| RT2_339.p1 | WSC domain-containing protein | Responses to stress cues and metal ions |
| RT2_12327.p1 | Xylosidase arabinofuranosidase | Involved in xylan degradation |
| RT2_15735.p1 | Zinc-binding dehydrogenase | Role in fungal infection |

**Table S7:** Uniquely expressed effector candidate genes during CAT fusion.

| Transcript_ID | Gene names | Possible functions |
|---|---|---|
| RT3_12055_m.41570 | ABC transporter transmembrane region | Involved in membrane transport |
| RT3_6147_m.31990 | Acetyl-CoA acetyltransferase | Role against oxidative and cell wall stresses |
| RT3_12912_m.42432 | Alkaline proteinase | Play an important role in fungal physiology and development |
| RT3_107_m.1859 | Alpha-glucan | Role in virulence |
| RT3_5369_m.29859 | Alpha-L-arabinofuranosidase B, catalytic | Involved in the degradation of arabinoxylan a major component of plant hemicellulose |
| RT3_2561_m.19389 | Alpha-L-fucosidase | Play important roles in several biological processes like signal transduction |
| RT3_780_m.8486 | alpha-L-rhamnosidase | Degradation of plant cell wall sugar L-rhamnose |
| RT3_4885_m.28425 | Amine oxidase | Role in cell differentiation, growth and signalling |
| RT3_1558_m.13860 | AMP-binding enzyme | It functions as a general fungal virulence factor in plant pathogenic ascomycetes |
| RT3_17141_m.45950 | Ankyrin-3-like protein 3 | Play key roles in activities such as cell motility, activation, proliferation, contact, and the maintenance of specialized membrane domains |
| RT3_6713_m.33305 | Arginase family | Role in host–fungus interaction |
| RT3_870_m.9203 | Aspartic endopeptidase | Role in cell-wall assembly, remodelling and cell wall integrity |
| RT3_16782_m.45667 | Beta-ig-h3 fasciclin | Play roles in fungal development; also triggers signaling pathways mediating adhesion and migration of vascular smooth muscle cells |
| RT3_2825_m.20686 | Blastomyces yeast-phase-specific protein | Unknown function |
| RT3_8169_m.36196 | C6 transcription factor | Involved in meiosis, stress response, pleiotropic drug resistance |
| RT3_8826_m.37297 | Calcineurin-like phosphoesterase | Involved in stress survival and fungal virulence |
| RT3_2348_m.18321 | Calcium influx-promoting protein ehs1 | Involved in maintaining cell wall integrity |
| RT3_13397_m.42919 | Calcium-related spray protein | Specific role in calcium stress response |
| RT3_1847_m.15557 | Catalase | Important for growth and the start of cell differentiation |

| | | |
|---|---|---|
| RT3_1557_m.13846 | Cellulosome enzyme | Degrades lignocellulose, cellulose and hemicellulose in plant cell wall |
| RT3_10144_m.39239 | Chitin synthesis regulation, resistance to Congo red | Plays vital roles against stress, noxious chemicals and osmotic pressure changes |
| RT3_4302_m.26566 | Chitinase-1 | Hydrolytic enzymes that break down glycosidic bonds in chitin |
| RT3_7671_m.35279 | Complex I intermediate-associated protein 30 | Increases susceptibility to oxidative stress and phleomycin |
| RT3_20705_m.48270 | Coronin | Role in the organization and dynamics of actin and F-actin remodeling |
| RT3_11624_m.41121 | Cupin superfamily (DUF985) | Function is unknown |
| RT3_12971_m.42487 | Cytochrome oxidase assembly protein | An integral protein of the inner mitochondrial membrane that is essential for cytochrome c oxidase assembly. Also have a role in the response to hydrogen peroxide exposure |
| RT3_100_m.1747 | Cytochrome p450 | Role in redox signaling |
| RT3_7169_m.34287 | Dihydrodipicolinate synthetase | Role in fungal development, pathogenesis and stress responses |
| RT3_4085_m.25779 | Dolichyl-diphosphooligosaccharide--protein glycosyltransferase subunit OST2 | Role in protein modification |
| RT3_306_m.4196 | Duf92 domain protein | Function is unknown |
| RT3_6853_m.33623 | Endo-beta-1,6-glucanase | Role in cell wall softening |
| RT3_8524_m.36827 | Enoyl-CoA hydratase/isomerase | Role in the biosynthesis of siderophore ferrichrome A which is contributing to pathogenicity |
| RT3_10427_m.39639 | Extracellular serine-rich protein | Perform a variety of functions e.g. protein maturation, signal peptide cleavage, signal transduction etc |
| RT3_156_m.2486 | FKBP-type peptidyl-prolyl cis-trans isomerase | Role in signaling. FKBP12s bind the immunosuppressive drug FK506 to inhibit the phosphatase calcineurin (CaN). CaN is required for virulence |
| RT3_13797_m.43265 | Frequency clock protein | Regulates various aspects of the circadian clock in *Neurospora crassa* |
| RT3_9389_m.38176 | Fructosyl amino acid | Role in conidial germination |
| RT3_4379_m.26842 | Fungal specific transcription | Essential regulators of gene expression in a cell, involved in meiosis, stress response, |

| | | |
|---|---|---|
| | factor domain | pleiotropic drug resistance etc. |
| RT3_16749_m.45643 | Galactoside-binding lectin | Role in fruiting body development |
| RT3_3953_m.25310 | Gamma-glutamyltranspeptidase | Role in the vacuolar transport and metabolism of glutathione |
| RT3_568_m.6708 | Glucoamylase | Belongs to glycosyl hydrolase family, role in plant cell wall degradation |
| RT3_19701_m.47620 | Glycoside hydrolase | Degrades cellulase, hemicellulose and lignin found in plant cell wall |
| RT3_2582_m.19489 | Glycosyl hydrolase family 3 N terminal domain-containing protein | Degrades cellulase, hemicellulose and lignin found in plant cell wall |
| RT3_11645_m.41138 | Glycosyl hydrolase family 45 | Degrades cellulase, hemicellulose and lignin found in plant cell wall |
| RT3_6645_m.33149 | Glycosyl hydrolase, family 43 | Degrades cellulase, hemicellulose and lignin found in plant cell wall |
| RT3_2827_m.20695 | Glycosyltransferase family 31 | Degrades cellulase, hemicellulose and lignin found in plant cell wall |
| RT3_861_m.9128 | Gpi anchored cell wall | Role in cell wall biogenesis and integrity |
| RT3_2987_m.21396 | GPI biosynthesis protein family Pig-F | Involved in GPI anchor biosynthesis and therefore, required for cell wall biogenesis and normal hyphal growth |
| RT3_25_m.564 | Hkr1p | Role in mitogen-activated protein kinase pathway |
| RT3_4431_m.27028 | Hypersensitive response-inducing protein | Triggers defense response |
| RT3_14213_m.43634 | Immunoglobulin I-set domain-containing protein | Are found in several cell adhesion molecules |
| RT3_7054_m.34066 | Inosine/uridine-preferring nucleoside hydrolase | Has hydrolase activity |
| RT3_18980_m.47151 | KR domain-containing protein | Role in xylan degradation |
| RT3_19664_m.47603 | Kynurenine 3-monooxygenase | Involved in the growth, development, and pathogenicity of *Botrytis cinerea* |
| RT3_5356_m.29823 | L-amino acid oxidase | Plays role in oxidative stress. |
| RT3_4165_m.26064 | L-ascorbate oxidase | Might be involved in plant pathogenesis, fungal development and signaling pathways under |

| | | |
|---|---|---|
| | | hypoxic condition |
| RT3_12332_m.41840 | LipA and NB-ARC domain-containing protein | May function as key integrators of stress and nutrient availability signals |
| RT3_3056_m.21708 | Long chronological lifespan protein 2 | Probable component of the endoplasmic reticulum-associated degradation (ERAD) pathway |
| RT3_4434_m.27038 | Malic enzyme | Role in fatty acid synthesis |
| RT3_5865_m.31259 | MATE efflux family protein | Expression of MATE family proteins might alters development, stress responses and pathogen susceptibility |
| RT3_2892_m.20978 | Mid2 like cell wall stress sensor | Required for cell wall integrity signalling pathway |
| RT3_18063_m.46545 | MIP family channel protein | Involved in the transport of water and neutral solutes across the membranes. |
| RT3_1783_m.15192 | Mixed-linked glucanase | Role in plant cell wall degradation |
| RT3_2110_m.17059 | ML domain-containing protein | Domain involved in innate immunity and lipid metabolism |
| RT3_268_m.3768 | NCS1 nucleoside transporter | NCS1 proteins are $H^+$ or $Na^+$ symporters responsible for the uptake of purines, pyrimidines or related metabolites |
| RT3_2176_m.17433 | Nitric oxide synthase | Role in the light-induced development of sporangiophores |
| RT3_8078_m.36041 | N-terminal domain on NACHT_NTPase and P-loop NTPases | Unknown function |
| RT3_15066_m.44317 | Nucleoside diphosphate kinase | Regulation of spore and sclerotia development |
| RT3_1557_m.13846 | O-Glycosyl hydrolase family 30 | Role in plant cell wall degradation |
| RT3_704_m.7811 | PAN domain | Essential for RasA-mediated morphogenetic signaling |
| RT3_12332_m.41840 | Pathogen effector; putative necrosis-inducing factor | Virulence factor |
| RT3_6968_m.33865 | Pectin lyase 2 | Degrades pectin of plant cell wall |

| | | |
|---|---|---|
| RT3_5718_m.30849 | Pectinesterase | Facilitate plant cell wall modification and subsequent breakdown |
| RT3_6809_m.33525 | Peptidase dimerisation domain | Role in pathogenesis |
| RT3_6809_m.33525 | Peptidase family M20/M25/M40 | Role in pathogenesis |
| RT3_156_m.2486 | Peptidylprolyl isomerase | The ppci1 gene plays important roles in growth, conidiation, and sclerotia formation |
| RT3_455_m.5677 | Phosphorylase superfamily | Affect a wide variety of biological processes, including morphology and virulence |
| RT3_11518_m.41005 | Platelet-activating factor acetylhydrolase | In the *Aspergillus*, it has been shown to control the migration of nuclei |
| RT3_18656_m.46944 | Polygalacturonase 3 | It is cell wall degrading enzyme |
| RT3_18612_m.46914 | PQ loop repeat | PQ-loop proteins might function as cargo receptors in vesicle transport |
| RT3_11944_m.41457 | Probable lipid transfer | The members of this family are probably involved in lipid transfer |
| RT3_21269_m.48652 | Protein kinase domain | Role in a multitude of cellular processes, including division, proliferation, apoptosis, and differentiation, cell wall integrity |
| RT3_21269_m.48652 | Protein serine/threonine kinase | Has many roles in cellular function like regulation of signaling pathways |
| RT3_3541_m.23752 | Protein YOP1 | Involved in membrane/vesicle trafficking |
| RT3_543_m.6478 | Proteolipid membrane potential modulator | Pmp3 is an evolutionarily conserved proteolipid in the plasma membrane which, in *S. pombe*, is transcriptionally regulated by the Spc1 stress MAPK (mitogen-activated protein kinases) pathway |
| RT3_571_m.6736 | Pyridoxamine 5'-phosphate oxidase | Role in oxidative stress |
| RT3_7114_m.34184 | Ribonuclease T2 family protein | Tomato T2 ribonuclease LE is involved in the response to pathogens |
| RT3_10423_m.39633 | RNA-binding domain-containing protein | RBPs play key roles in post-transcriptional processes |
| RT3_13063_m.42567 | RTA1 like protein | Expression is induced under both low-heme and low-oxygen conditions |
| RT3_187_m.2871 | Signal peptidase subunit 3 | SPC3 is required for signal peptidase activity and is essential for viability |

| | | |
|---|---|---|
| RT3_2326_m.18214 | SOCE-associated regulatory factor of calcium homoeostasis | Negative regulator of intracellular calcium signaling and calcium transport play a role in growth, virulence, and stress resistance |
| RT3_9184_m.37861 | Sodium/hydrogen exchanger family protein | Is a membrane transport protein involved in signaling |
| RT3_15296_m.44520 | Spherulation-specific family 4 | Associated with formation of spherules or spores under starvation conditions |
| RT3_2348_m.18321 | Stretch-activated Ca2+-permeable channel component | MID1 is a yeast Saccharomyces cerevisiae gene encoding a plasma membrane protein required for $Ca^{2+}$ influx induced by the mating pheromone, alpha-factor |
| RT3_2386_m.18522 | SWIM zinc finger | Is a zinc finger protein |
| RT3_13567_m.43062 | Tetraspanin | Pls1, which is required for pathogenicity and Tsp2, whose function is unknown. Tetraspanins are a superfamily of small integral membrane proteins |
| RT3_938_m.9738 | Transient receptor potential ion channel | Exhibiting activation by hyperosmolarity, temperature increase, cytosolic $Ca^{2+}$ elevation, membrane potential, and $H^2O^2$ application |
| RT3_18634_m.46933 | Ulp1 protease family protein | Has an essential role in the G2/M phase of the cell cycle |
| RT3_1_m.4 | WD40 domain-containing protein | A WD40 repeat protein regulates fungal cell differentiation |
| RT3_3897_m.25109 | Wlm domain-containing protein (Fragment) | It is related to the DNA damage response proteins WSS1 involved in sister chromatid separation and segregation |
| RT3_10404_m.39608 | WSC domain-containing protein | Responses to stress cues and metal ions |

**Table S8:** List of primers used in the study for real time qPCR validation.

| Transcript_ID | Primer name | Primer sequences | Tm (˚C) |
|---|---|---|---|
| β-tubulin gene | Btubulin1F | TCCCGAACAATGTGCAGACA | 59 |
| | Btubulin1R | AGAACGCCTTTCTGCGAAAC | 58 |
| RT1_3180 | RT1GLYCOF | GGTAAACGTCGGAGTCGGTA | 60 |
| | RT1GLYCOR | GAGTCCATCGTCCTCCTGAA | 60 |
| RT2_4748 | RT1MFS1F | AGACCCATGAGGATGGTCTG | 60 |
| | RT1MFS1R | CTCGATGTCCTCAACAAGCA | 60 |
| RT1_4882 | RT1CBFF | TCTCCTGAAGCTGTGAAGCA | 60 |
| | RT1CBFR | AATTCCCCCATTCACCTTTC | 60 |
| RT1_48003 | RT1STEF | CATCATGTTGGCCAGCTACC | 61 |
| | RT1STER | AGGAGGCGTAGATCTGCTTG | 60 |
| RT1_47185 | RT1PPGF | GGAATCACCCAGGTTACGAA | 60 |
| | RT1PPGR | ATTGCTGTGTGTTTGGATGC | 60 |
| RT1_12359 | RT1ALF | TGCAGAAGTCGTAGGCAATG | 60 |
| | RT1ALR | GAACAACCTGCAGAGCATCA | 60 |
| RT3_2628 | RT1DTPF | ACGGTTGACCGGTATGAGAG | 60 |
| | RT1DTPR | GCCCCAGTGAATGAAGTGTT | 60 |
| RT3_7259 | RT1RTAF | TTACCGTCGGTGGCTTATTC | 60 |
| | RT1RTAR | ATGTTCGGCCTTGACAGTTC | 60 |
| RT3_2068 | RT1FEBF | TGCTTCTGGCCGTAGAAGTT | 60 |
| | RT1FEBR | GGTCTCAGAGGGGAATGTCA | 60 |
| RT1_45519 | RT1SET9F | GCCTCCTACGACGACTTTCT | 60 |
| | RT1SET9R | GGGGATTGTGGTCCAGTAAA | 60 |
| RT1_1703 | RT1LARAF | CCACATGGAGGCCATCTACT | 60 |
| | RT1LARAR | GTAGTGAAGCGGGTGGTGAT | 60 |
| RT1_3153 | RT1HETF | TGATCACTAGGCAGCAGTGG | 60 |
| | RT1HETR | CACATCGAGAGTGCTTGGAA | 60 |
| RT2_5451 | RT2PECT1F | CGACGAGGTAGTCCTTGAGC | 60 |
| | RT2PECT1R | GGCCTACTCCAACAGCTTCA | 60 |
| RT2_6816 | RT2CBMF | GGTGTTTGTTGGGTGAGGTT | 60 |
| | RT2CBMR | TCCTCATCCGGTTCTTCATC | 60 |
| RT2_11038 | RT2FGGYF | GCGTTGATCTTCTCCGTCTC | 60 |
| | RT2FGGYR | CTGCCAAGATCAAGGGAATC | 60 |
| RT2_9885 | RT2HRIF | TTTACAGCCCTGGACTGAGC | 60 |
| | RT2HRIR | ACCTCGAGTAAGCGGTTTGA | 60 |
| RT2_3495 | RT2PDAF | AAGACGACTCCACGGGTATG | 60 |
| | RT2PDAR | GGTGGACGGAAGTAGTTGGA | 60 |

| | | | |
|---|---|---|---|
| RT2_7978 | RT2MPF | CTGAGGAGGCCTACAACTGC | 60 |
| | RT2MPR | GGTGTCCCAGTAGGAGTGGA | 60 |
| RT2_20649 | RT3MFSF | AGAGCCGATGCAGACAGAAT | 60 |
| | RT3MFSR | ATGGCTTTCGAACAATTTGG | 60 |
| RT3_7864 | RT3SMF | AACATCTCCGCAAACCAATC | 60 |
| | RT3SMR | TGGAGGATGAGGAGGAGAGA | 60 |
| RT3_14378 | RT3FDF | GTTTCGAAGTCAGCCCAGAA | 60 |
| | RT3FDR | CTCGCCTGCAGAGAGCTAGT | 60 |
| RT1_35493 | RT3ZNF | CAACCGTTGGTTTGGGATAC | 60 |
| | RT3ZNR | ATGAAGATCCTGCCAGTCGT | 60 |
| RT1_43231 | RT3DUFF | CCGATGTAGACGGTGTTTCC | 60 |
| | RT3DUFR | ATCTCCATCTGGGGCTCTTT | 60 |
| RT1_51163 | RT3SHEF | TTGGCGAGAGCGACACAG | 62 |
| | RT3SHER | ACGAAGGTTGGTGTCATTGTG | 60 |
| RT1_50197 | RT3DLPAF | TGACGTCAACATCCGACATC | 60 |
| | RT3DLPAR | CCTCTAGGAAGCCTCCGTCA | 60 |
| RT3_54854 | RT3BAXF | GGGATGTAGCTGCGTATCGT | 60 |
| | RT3BAXR | CTGAGCTTCGGCACCATGTA | 60 |
| RT1_23825 | RT3SRDF | ATTGGCCTTCTCACACAACC | 60 |
| | RT3SRDR | CCGTCCTGGGTGTTATTTGT | 60 |
